# ApoE-Dependent Lipid Handling by Median Eminence Microglia Preserves Myelin Integrity and Metabolic Function

**DOI:** 10.64898/2026.02.18.706643

**Authors:** ER McGrath, A Folick, LJ Morrissette, SM Brown Mayfield, V Pham, SN Pillutla, MK Choi, RT Cheang, WR Bolus, A Schwendeman, V Ntranos, SK Koliwad, M Valdearcos

## Abstract

Microglia regulate hypothalamic control of systemic metabolism, but the mechanisms underlying their contribution remain unclear. Here, we identify a distinct apolipoprotein E (*ApoE*)⁺ microglial population enriched in the median eminence (ME), a brain region involved in sensing peripheral cues and metabolic regulation. These microglia engage multiple functional programs related to lipid handling, interferon signaling, and stress responses that are differentially regulated within the ME. Consumption of a Western diet (WD) increased interferon signaling and lipid accumulation in ME microglia. Expression of the human *APOE4* isoform in mice exacerbated microglial lipid dysregulation, interferon signaling, and impaired ME myelin organization. Deleting *APOE* in microglia attenuated their ability to couple lipid accumulation to interferon signaling, identifying microglial APOE as a cell-intrinsic determinant of interferon responses. Finally, selective activation of liver X receptor signaling using synthetic HDL nanoparticles restored microglial lipid homeostasis, improved hypothalamic leptin responsiveness, and limited weight gain in WD–fed mice. Together, these findings define an *Apoe*-dependent regulatory program in ME microglia that is therapeutically targetable and clarify how nutritional stress disrupts hypothalamic control of metabolic homeostasis.

## Introduction

Microglia, the resident macrophages of the central nervous system, are increasingly recognized as metabolically active cells with roles far beyond their canonical immune-surveillance functions. In addition to shaping inflammatory tone, microglia regulate lipid handling, phagocytosis, and coordinated intracellular signaling to enable adaptation to metabolic and environmental stress. Single-cell transcriptomic studies have revealed substantial heterogeneity among microglia across brain regions, developmental stages, and disease contexts, challenging the notion of a uniform microglial response to stimuli^1–3^. Distinct transcriptional programs, including disease-associated microglia (DAM)^4^, interferon-responsive microglia (IRM)^5^, and white matter–associated microglia (WAM)^6^ illustrate how microglia adopt specialized states in response to metabolic, inflammatory, or degenerative stress. In demyelinating and aging contexts, microglia upregulate *Itgax* (CD11c) together with Clec7a, a pattern-recognition receptor associated with phagocytic and lipid-handling functions, defining adaptations to lipid-rich microenvironments reflective of myelin engagement^6^. Genetically targeting these states shows that Clec7a^+^ microglia are dynamic and context-dependent, and perturbing their adaptive ability alters disease progression and recovery in demyelination models^7^. Together, these findings highlight the flexibility of microglia to remodel their metabolic programs in response to tissue-specific demands.

These metabolic functions could be particularly relevant in the hypothalamus, a central regulator of systemic energy and glucose homeostasis. Within the hypothalamus, the median eminence (ME) is a specialized circumventricular region characterized by fenestrated vasculature and specific tanycytic barriers, enabling dynamic access of circulating hormones and nutrients to adjacent circuits in the hypothalamic arcuate nucleus (ARC)^8–10^. Thus, ME-resident microglia are positioned at the front line of metabolic signaling, where they must integrate fluctuating peripheral cues while preserving tissue integrity under conditions of nutritional stress. Consistent with this role, emerging evidence indicates that microglia actively shape hypothalamic circuits controlling energy and glucose homeostasis, including by regulating inflammatory tone, barrier properties, and neuron-glia interactions^11–14^. Despite growing appreciation of microglial involvement in hypothalamic physiology, the cellular programs that enable microglia to maintain lipid and metabolic homeostasis in this unique niche are poorly understood.

Recent work has revealed that the ME myelin undergoes continuous turnover, establishing this region as a rare site of active myelin remodeling outside of development or overt injury^15^. This plasticity likely reflects the ME’s role as a structural and functional gateway between the periphery and the brain, as well as its exposure to circulating metabolic signals^8,16^. Sustained myelin remodeling requires efficient lipid clearance and recycling, coordinated through interactions between oligodendrocytes, which generate new myelin, and microglia, which phagocytose myelin debris and recycle cholesterol^17,18^. Consistent with this requirement, microglia-deficient mouse models have profound defects in myelination and white matter organization^19^. Given that myelin represents the brain’s largest cholesterol reservoir, disrupting microglial lipid recycling limits cholesterol availability for myelin synthesis, leading to myelin fragmentation and associated inflammatory stress^20^. Importantly, perturbing oligodendrocyte lineage cells in the ME impairs leptin sensitivity and promotes metabolic dysregulation, highlighting the importance of myelin integrity for hypothalamic metabolic control^21^. Together, these observations identify myelin remodeling as a dynamic, homeostatic feature of the ME and raise the question of how microglial lipid metabolism supports this process during nutritional challenges.

Cholesterol homeostasis in the brain relies on tightly regulated local recycling mechanisms that are distinct from those in peripheral tissues^22,23^. Because cholesterol cannot be enzymatically degraded and exchange across the blood–brain barrier is limited, excess cellular cholesterol is packaged and redistributed between brain cells via cellular efflux pathways^23,24^. Microglia are central to this process by exporting cholesterol in apolipoprotein E (ApoE)–containing lipoproteins through a pathway governed by signaling via liver X receptor (LXR), a master transcriptional regulator of cellular cholesterol homeostasis^25^. Disrupting this pathway leads to the emergence of lipid droplet–accumulating microglia (LDAM), characterized by impaired cholesterol efflux, defective phagocytic capacity, and maladaptive inflammatory signaling^26^. LDAM have been described in aging and neurodegenerative contexts, where inadequate microglial cholesterol handling is associated with oxidative stress and functional decline^26^. Notably, chronic consumption of a Western diet (WD) suppresses LXR-driven cholesterol efflux in microglia, impairing myelin debris clearance and regenerative responses following demyelination^27^. Whether similar lipid-laden microglial states emerge in the hypothalamus during nutritional excess and how they impact myelin integrity and metabolic signaling remain unclear.

In contrast to murine ApoE, human APOE, the principal apolipoprotein in the human brain^28^, exists in three common isoforms (APOE2, APOE3, APOE4) that differ in lipid-binding properties, receptor interactions, and inflammatory tone^29,30^. Compared to *APOE3*, the predominant human allele, myeloid cells expressing *APOE4* have a reduced capacity to support cholesterol efflux and are associated with heightened inflammatory signaling^25,31^. Importantly, the development of isoform-specific humanized *APOE* models has allowed investigators to interrogate how *APOE* genotype shapes microglial lipid handling and stress responses in vivo^32^. However, whether *APOE* isoforms differentially influence microglial responses to nutritional excess in metabolically sensitive hypothalamic niches is unknown.

In this study, we identify a transcriptionally distinct *Apoe*⁺ microglial population enriched in the ME that encompasses multiple functional programs supporting lipid handling and myelin organization at this neurovascular interface. We show that WD feeding shifts microglial programs toward lipid accumulation and stress-associated interferon signaling, while expression of the human *APOE4* isoform exacerbates microglial lipid dysregulation and vulnerability of myelin organization. Using genetic and pharmacological approaches, we further demonstrate that diet-induced shifts in microglial lipid-handling programs are reversible, and targeted activation of LXR signaling in microglia using synthetic HDL nanopa DAPI eventsrticles is sufficient to restore microglial lipid homeostasis and improve metabolic responses to dietary excess.

## Results

### Identification of a distinct Apoe^+^ microglial population enriched in the hypothalamic ME of adult mice

Microglia are underrepresented in most published hypothalamic single-cell datasets, which typically prioritize broad cellular diversity and rely on dissociation protocols that compromise the survival of specialized immune populations. To overcome this limitation, we optimized a dissociation and enrichment strategy that maximizes recovery of viable microglia while minimizing transcriptional stress, enabling a single-cell RNA sequencing (scRNA-seq) workflow that captures the transcriptional heterogeneity of hypothalamic microglia.

Using this approach, microglia were isolated from the mediobasal hypothalamus (MBH) of mice fed either a low-fat, low-sugar control diet (CD) or a high-fat, high-sugar WD. Cells were enriched by fluorescence-activated cell sorting (FACS) of CD11b^+^ DAPI^−^ events, barcode-labeled, multiplexed, and processed in parallel across dietary conditions (**Figure 1A**). This strategy excluded debris, dead cells, and doublets, yielding robust enrichment of viable myeloid cells (**Figure S1**). The MULTI-seq lipid-conjugated DNA barcoding^33^ strategy we used enabled pooling of biological replicates while minimizing batch effects, with high barcode assignment efficiency and minimal cross-contamination. Only one of ten samples was excluded due to insufficient barcode recovery, validating the reliability of the multiplexing approach (**Figure S2**).

**Figure 1.**
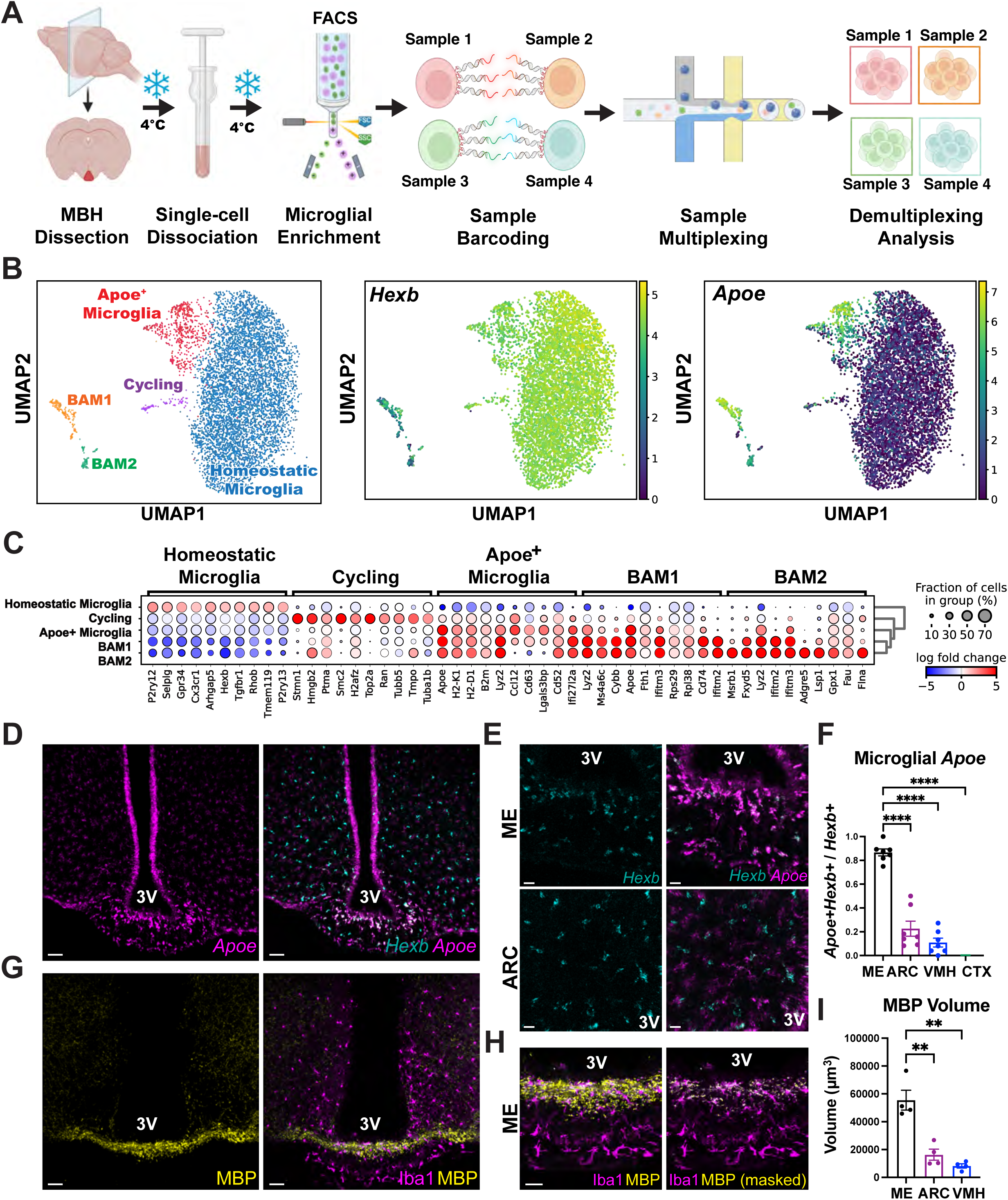
Identification of Apoe⁺ microglia engaged in myelin remodeling in the median eminence. (A) Schematic of MULTI-seq sample preparation and experimental workflow. Adult male mice were subjected to dietary manipulation for 4 weeks prior to mediobasal hypothalamus (MBH) dissection, FACS isolation of CD11b⁺DAPI⁻ cells, sample barcoding, multiplexing, and single-cell RNA sequencing. (B) UMAP visualization of MBH myeloid populations with feature plots showing normalized expression of *Hexb* and *Apoe*, identifying a distinct Apoe⁺ microglial population. (C) Dot plot showing expression of representative marker genes across transcriptionally defined microglial populations, including homeostatic, cycling, Apoe⁺ microglia, and border-associated macrophage (BAM) populations. Dot size indicates the fraction of cells expressing each gene, and color intensity reflects scaled average expression (blue to red, low to high). (D) Representative RNAscope images showing *Apoe* transcripts and *Apoe/Hexb* transcript colocalization in the MBH (scale bar, 50 µm). (E) Representative images showing co-localization of *Hexb* and *Apoe* transcripts in the ME and arcuate nucleus (ARC) (scale bar, 20 µm). (F) Quantification of the fraction of *Apoe*⁺ *Hexb*⁺ microglia across brain regions shown in (D) (n = 7; ****p < 0.0001). (G) Representative MBH images stained for myelin basic protein (MBP) and the microglial marker Iba1 (scale bar, 50 µm). (H) Higher-magnification images showing intracellular MBP signal within Iba1⁺ microglia in the ME (scale bar, 20 µm). (I) Quantification of the proportion of Iba1⁺ microglia containing intracellular MBP signal in the ME (n = 3; **p < 0.01). Data are presented as mean ± SEM and were analyzed using unpaired two-tailed *t* tests. 3V, third ventricle.

After quality control filtering, unsupervised clustering resolved several transcriptionally distinct myeloid populations, including transcriptional programs reflecting homeostatic microglia, which expressed canonical markers such as *P2ry12* and *Tmem119*, cycling microglia characterized by cell-cycle genes (*Stmn1*, *Hmgb2*, and *Top2a*), and two populations of border-associated macrophages (BAM1 and BAM2) with gene expression profiles distinct from resident microglia (**Figures 1B-C**). Notably, we identified a discrete cluster of microglia expressing high levels of the gene encoding ApoE (*Apoe*), along with genes associated with lysosomal and phagocytic activity (*Cd63*, *Lyz2*) and antigen presentation (*H2-K1*, *H2-D1*, and *B2m)*. Co-expression of *Hexb* confirmed that these *Apoe^+^* cells were bona fide microglia rather than infiltrating macrophages (**Figure 1B**). We refer to this microglial population as MG-Apoe.

To determine whether MG-Apoe occupy a specific anatomical niche within the MBH, we performed spatial analysis using *in situ* hybridization for *Apoe* and *Hexb*. Representative images are shown from mice fed a WD for 4 weeks to enhance the detection of this spatially restricted population. While *Hexb*^+^ microglia were broadly distributed across MBH subregions, MG-Apoe cells were strikingly enriched within the ME (**Figure 1D**). Indeed, high-magnification imaging revealed robust co-localization of *Apoe* and *Hexb* within the ME, whereas such co-localization was minimal in the adjacent ARC, despite comparable microglial density (**Figure 1E**), and quantification across hypothalamic and extrahypothalamic regions confirmed a selective enrichment of MG-Apoe cells in the ME relative to the ARC, ventromedial hypothalamus (VMH), and cerebral cortex (CTX) (**Figure 1F**). The ME is characterized by dense bundles of myelinated axons, prompting us to examine the spatial relationship between MG-Apoe microglia and local myelin. Immunohistochemical staining for myelin basic protein (MBP) and the microglial marker Iba1 revealed that ME microglia are positioned within myelin-rich zones (**Figure 1G**), with high-resolution imaging further revealing that this MBP signal is present within Iba1^+^ microglia in the ME (**Figure 1H**). Moreover, and consistent with these observations, ME microglia contained significantly greater MBP volume than did microglia in the ARC or VMH (**Figure 1I**). Thus, MG-Apoe microglia are a specifically myelin-associated population of microglia that are spatially restricted within the ME.

### MG-Apoe microglia in the ME comprise multiple transcriptionally distinct programs

Analysis of *Apoe* expression revealed substantial heterogeneity within the MG*-*Apoe population, prompting further resolution of transcriptional diversity within this population. Subclustering of MG*-*Apoe microglia resolved three distinct transcriptional programs, designated MG-Apoe^hi^, MG-Apoe^int,^ and MG-Apoe^lo^, based on relative *Apoe* expression levels, with each subcluster occupying discrete regions in the UMAP space (**Figure 2A**). Violin plots analysis confirmed that *Apoe* expression was indeed graded across these states in descending order from MG-Apoe^hi^ to MG-Apoe^int,^ and then to MG-Apoe^lo^, with expression in MG-Apoe^lo^ remaining higher than in homeostatic microglia (**Figure 2B**). We compared the profiles of each MG-Apoe transcriptional program to established microglial programs associated with aging and disease, including disease-associated microglia (DAM), interferon-responsive microglia (IRM), and a senescence-associated gene program (SenMayo). Scoring against this reference set of signatures revealed distinct, non-overlapping associations across MG-Apoe states, suggesting that these states reflect multiple engagement of partially independent response modules rather than a single uniform transcriptional mode of activation (**Figures 2C-E; Figure S3**). MG-Apoe^int^ was preferentially enriched for the IRM signature, whereas MG-Apoe^lo^ had the highest SenMayo score, consistent with a stress- and senescence-associated transcriptional profile. MG-Apoe^hi^ showed intermediate enrichment across programs, suggesting a mixed or potentially transitional state.

**Figure 2.**
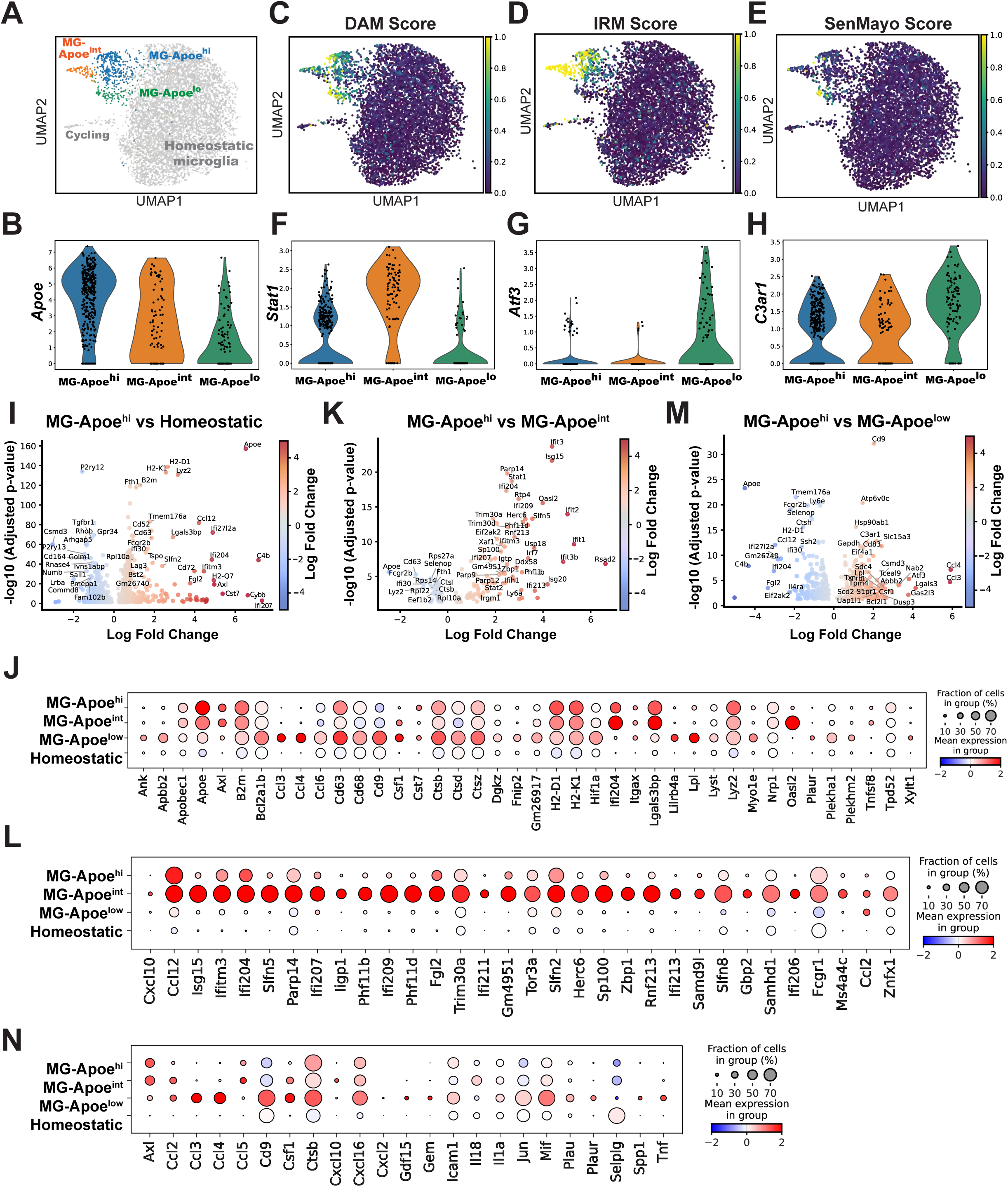
Apoe⁺ microglia comprise three transcriptionally distinct programs in the median eminence. (A) UMAP visualization of MG-Apoe microglia, subclustered into three transcriptionally distinct states (MG-Apoe^hi^, MG-Apoe^int^, and MG-Apoe^lo^), shown relative to homeostatic and cycling microglia. (B) Violin plot showing graded *Apoe* expression across MG-Apoe subclusters, with the highest expression in MG-Apoe^hi^ and progressively lower expression in MG-Apoe^int^ and MG-Apoe^lo^. (C–E) UMAP feature plots showing enrichment scores for disease-associated microglia (DAM) (C), interferon-responsive microglia (IRM) (D), and the SenMayo senescence signature (E), demonstrating preferential enrichment of IRM programs in MG-Apoe^int^ and senescence-associated programs in MG-Apoe^lo^. (F) Violin plot showing selective enrichment of *Stat1* expression in MG-Apoe^int^. (G) Violin plot showing enrichment of the stress-responsive transcription factor *Atf3* in MG-Apoe^lo^. (H) Violin plot showing selective enrichment of *C3ar1* mRNA in MG-Apoe^lo^. (I) Volcano plot showing differential gene expression between MG-Apoe^hi^ and homeostatic microglia. (J) Dot plot visualization of DAM signature genes across MG-Apoe subclusters and homeostatic microglia. (K) Volcano plot showing differential gene expression between MG-Apoe^hi^ and MG-Apoe^int^. (L) Dot plot visualization of IRM signature genes across MG-Apoe subclusters and homeostatic microglia. (M) Volcano plot showing differential gene expression between MG-Apoe^hi^ and MG-Apoe^lo^. (N) Dot plot visualization of SenMayo senescence-associated genes across MG-Apoe subclusters and homeostatic microglia. Dot sizes in panels (J), (L), and (N) represent the fraction of cells expressing each gene, and color intensity reflects scaled average expression (blue to red, low to high).

Differential expression analysis identified molecular features distinguishing each subcluster. *Stat1* emerged as a defining marker of the IRM-enriched MG-Apoe^int^ population (**Figure 2F**). In contrast, MG-Apoe^lo^ microglia selectively expressed stress- and lipid-associated genes, including the transcription factor *Atf3*^34,35^ (**Figure 2G**) and the complement receptor *C3ar1* (**Figure 2H**), which have been implicated in myelin lipid handling and lipid droplet (LD) formation^36^. Comparative analyses revealed that MG-Apoe^hi^ microglia upregulated immune and antigen presentation pathways with coordinated expression of DAM-associated transcripts while downregulating canonical homeostatic markers (**Figure 2I**), distinguishing this population from homeostatic microglia (**Figure 2J**). Using MG-Apoe^hi^ as a reference state, direct comparison revealed selective upregulation of interferon-responsive genes in MG-Apoe^int^ microglia (**Figure 2K**), consistent with its preferential enrichment for IRM signatures (**Figure 2L**). In contrast, MG-Apoe^lo^ microglia had increased expression of genes involved in lipid metabolism, stress responses, and chemokine signaling, including *Cd9*, *Lpl*, *C3ar1*, and *Atf3* (**Figure 2M**). Extending this analysis across all microglia populations confirmed preferential enrichment of inflammatory and stress-associated transcripts in MG-Apoe^lo^ relative to other MG-Apoe states and homeostatic microglia (**Figure 2N**). Together, these findings reveal that MG-Apoe microglia in the median eminence comprise multiple transcriptional states spanning interferon-responsive, immune-activated, and metabolically stressed programs.

### MG-Apoe in the ME are defined by a myelin-dependent CD11c⁺ program and a myelin-independent interferon-associated program

To visualize MG-Apoe states within intact hypothalamic tissue, we used CD11c (*Itgax*) expression as a surrogate marker, as it was selectively enriched across all three MG-Apoe subclusters but absent from homeostatic microglia in our single-cell dataset (**Figure S4A–B**). Consistent with this designation, high-resolution imaging revealed that CD11c⁺ microglia were selectively localized to the ME and largely absent from adjacent hypothalamic regions (**Figure S4C**). Quantitative analysis further confirmed significant enrichment of CD11c signal in the ME relative to the ARC and VMH, paralleling regional differences in MBP abundance (**Figure S4D–E**), and thereby positioning CD11c^+^ microglia within the myelin-rich niche of the ME. Indeed, CD11c⁺ microglia in the ME had higher expression levels of both the lysosomal marker CD68 and the interferon-associated transcription factor STAT1 compared with microglia in neighboring regions (**Figure S4F–H**), consistent with engagement of lipid-handling and stress-responsive programs. Moreover, Clec7a expression, a receptor previously linked to microglial responses to myelin remodeling and debris clearance in models of demyelination and neurodegeneration^7,37^, co-localized extensively with CD11c in ME microglia (**Figure S4I**), indicating that these markers identify the same myelin-associated CD11c⁺ MG-Apoe population.

To test whether these microglial programs differ in their dependence on ongoing myelin production, we conditionally disrupted oligodendrocyte myelination by deleting myelin regulatory factor (*Myrf*) in the oligodendrocyte lineage using *Pdgfrα*-Cre^ER^ mice as previously described^15^. This manipulation markedly reduced MBP immunoreactivity in the ME (**Figure S5A-B**) and decreased MBP signal within Iba1^+^ microglia (**Figure S5C**). Whereas disrupting myelin production selectively attenuated microglial Clec7a expression in the ME of mice fed a WD, it had no effect on microglial interferon-associated STAT1 response (**Figure S5D-F**). Together, these findings distinguish two diet-induced lipid-handling microglial programs in the ME: a myelin-dependent Clec7a⁺ program and a myelin-independent interferon-associated STAT1^+^ program.

### Chronic WD consumption concurrently stimulates interferon signaling and lipid accumulation in ME microglia

While scRNA-seq resolved distinct MG-Apoe⁺ transcriptional microglial programs among MBH microglia, this approach did not reveal significant changes in the relative abundance of these states following WD feeding. Consistent with this, the proportion of microglia assigned to each transcriptional cluster was comparable between CD and WD-fed mice (**Figure S2B**), and microglia from both dietary conditions occupied similar regions in the UMAP space (**Figure S2C**). These findings indicate that WD consumption does not substantially alter the representation of the MG-Apoe cluster, as resolved by scRNA-seq. However, the absence of detectable changes in cluster abundance does not exclude the presence of diet-specific functional response programs within these MG-Apoe states. Moreover, because scRNA-seq fundamentally does not capture changes in protein expression or cellular lipid accumulation, we next used immunohistochemical approaches to assess diet-induced microglial responses in tissue.

WD consumption induced robust upregulation of STAT1 in Iba1^+^ ME microglia after both 4 and 8 weeks (**Figure 3A-B**). In parallel, WD feeding markedly expanded the population of CD11c⁺ microglia in the ME, with progressive increases in both density and abundance over time (**Figure 3C-D**). Given the enrichment of lipid-handling transcriptional programs within *MG-*Apoe cells and exposure of the ME to peripheral metabolic cues, we next examined LD accumulation in this hypothalamic region. WD feeding substantially increased LD content within the ME, reflected by an increase in both total LD volume (**Figure 3E**) as well as LD accumulation specifically in microglia, as quantified by LD burden per Iba1^+^cell (**Figure 3F**). Notably, although LDs were detected across ME microglia in general, they were preferentially enriched in CD11c^+^ microglia, which we had found to be myelin-responsive (**Figure 3G**) and STAT1^+^ microglia, which we had found to be induced independent of myelin production (**Figure 3H**). Together, these data show that WD feeding concurrently induces interferon signaling and lipid accumulation within ME microglia, particularly among CD11c^+^ and STAT1^+^ cells, but that the diet-induced induction of lipid storage by these microglia is not exclusively coupled to myelin engagement.

**Figure 3.**
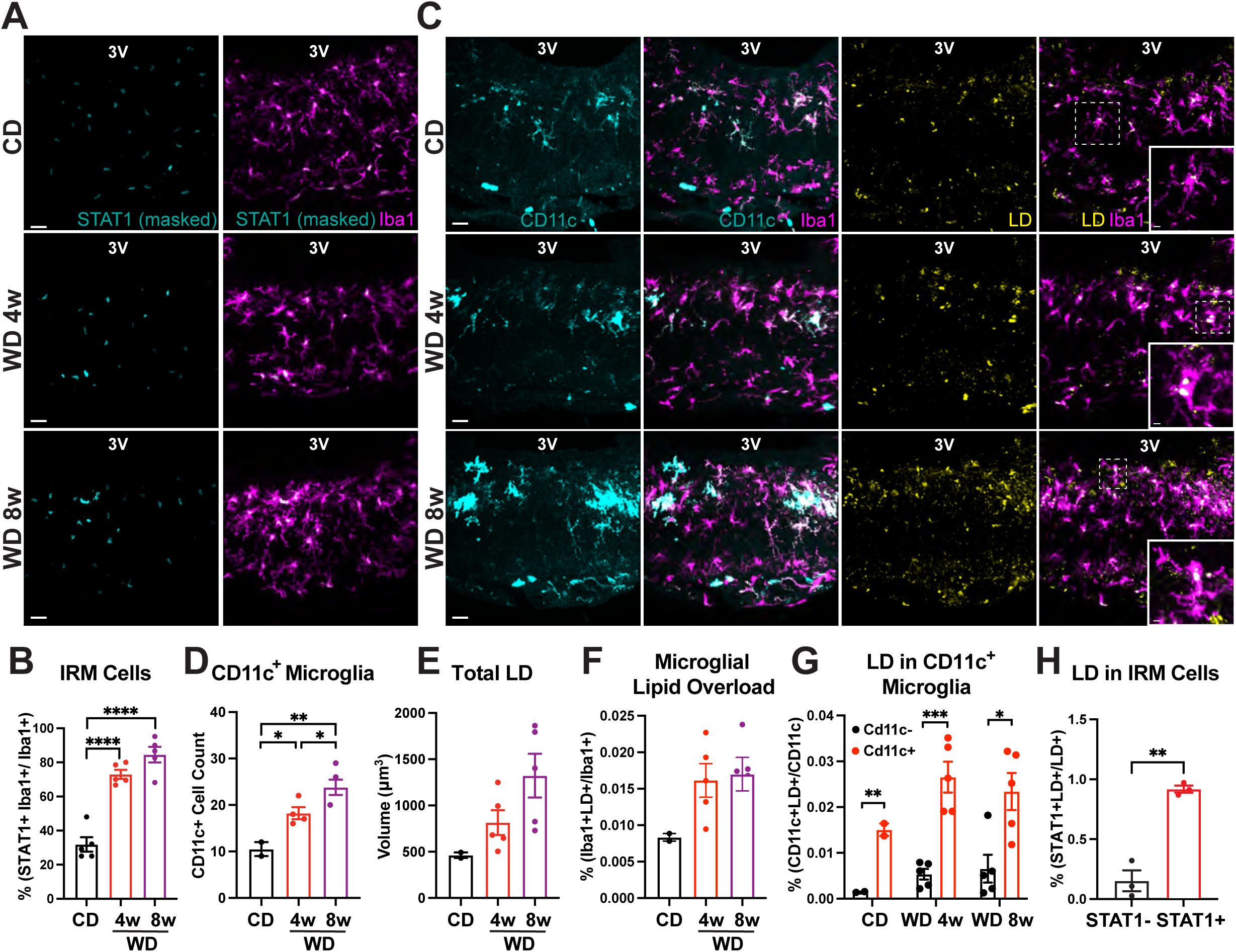
Consumption of a Western diet induces interferon signaling and microglial lipid accumulation in the median eminence. (A) Representative images showing STAT1 immunoreactivity within Iba1⁺ microglia in the median eminence (ME) of adult male wild-type mice fed a control diet (CD) or western diet (WD) for 4 or 8 weeks. STAT1 signal was quantified by sequential masking to DAPI and Iba1 to restrict analysis to microglial nuclei. (B) Quantification of STAT1 signal intensity in Iba1⁺ microglia shown in panel A (n = 5 mice per group; ****p < 0.0001). (C) Representative images of CD11c immunostaining and LipidSpot staining (used to label lipid droplets, LD) in Iba1⁺ microglia within the ME of mice fed a CD or WD for 4 or 8 weeks. (D–H) Quantification of microglial CD11c expression and lipid droplet accumulation shown in panels B and C (n = 2–5 mice per group; *p < 0.05, **p < 0.01, ***p < 0.001). Data were analyzed using unpaired two-tailed t-tests and are presented as mean ± SEM. Scale bars, 20 µm (insets, 5 µm). 3V, third ventricle.

### ApoE and Trem2 play distinct roles in lipid handling and interferon signaling in ME microglia under dietary stress

The ApoE-Trem2 pathway has been shown to regulate myeloid lipid sensing and state transitions, including the emergence of ApoE-dependent DAM states^6,38^. In other brain regions, activation of this pathway promotes LD accumulation in microglia in a Trem2-dependent manner following demyelinating injury^39,40^. While ApoE is broadly expressed across glial populations, Trem2 expression is largely restricted to myeloid cells, including microglia in the central nervous system and lipid-associated macrophages in peripheral metabolic tissues^41^. We therefore asked whether this pathway contributes to microglial responses in the ME during WD feeding and whether ApoE and Trem2 play distinct roles in shaping lipid-driven adaptations in this niche. WD feeding increased Trem2 expression in ME microglia (**Figure S6**), consistent with engagement of lipid-responsive pathways in this region. To assess the requirement of ApoE and Trem2 in microglial interactions with myelin, we compared ME microglia from *Apoe*-deficient (*Apoe*^-/-^ and *Trem2*-deficient (*Trem2*^-/-^) mice fed a WD with their respective wild-type counterparts under WD feeding. Notably, ME microglia from both *Apoe*^−/–^ and *Trem2*^-/-^ mice exhibited reduced Clec7a expression (**Figures 4A–B**), and showed diminished association with myelin, as reflected by reduced MBP signal within microglia and a lower myelin continuity index in the ME (**Figures 4C–D**). Together, these findings indicate that both ApoE and Trem2 are required for effective microglial engagement with myelin in the ME during dietary excess.

**Figure 4.**
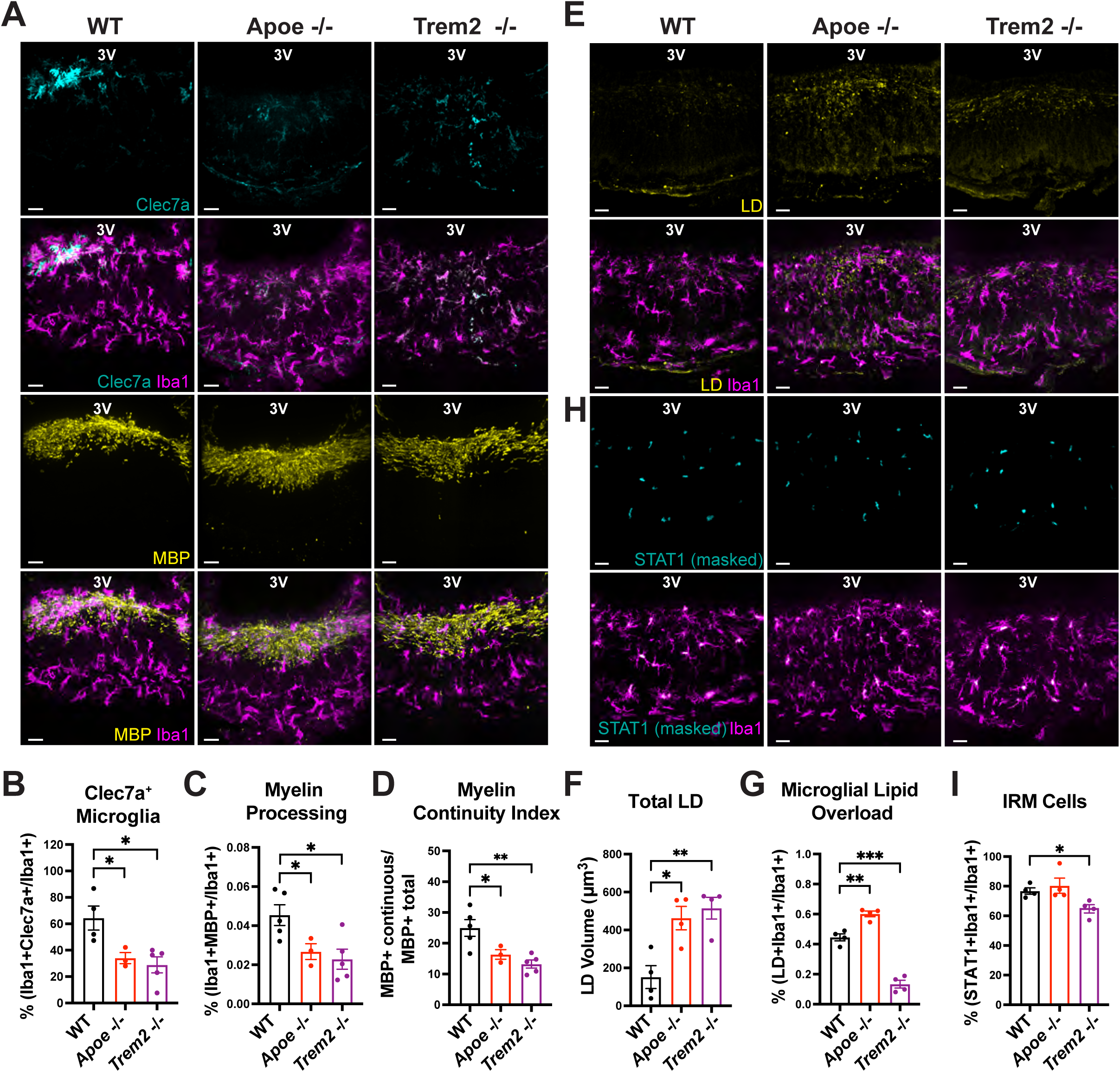
Loss of Apoe and Trem2 differentially alters lipid handling and interferon signaling in median eminence microglia during Western diet feeding. (A) Representative images of Clec7a or myelin basic protein (MBP) staining in Iba1⁺ microglia within the median eminence (ME) of adult male wild type (WT), *Apoe*⁻/⁻, and *Trem2*⁻/⁻ mice fed a western diet (WD) for 4 weeks. (B–D) Quantification of microglial Clec7a expression and microglia–myelin association from the dataset shown in panel A (n = 3–5 mice per group; *p < 0.05, **p < 0.01). (E) Representative images of LipidSpot staining used to label lipid droplets (LDs) in ME microglia. (F–G) Quantification of total LD volume and microglial LD accumulation from the dataset shown in panel E (n = 4 mice per group; *p < 0.05, **p < 0.01, ***p < 0.001). (H) Representative images showing STAT1 immunoreactivity within Iba1⁺ microglia in the ME of WT, Apoe⁻/⁻, and Trem2⁻/⁻ mice fed a WD for 4 weeks. STAT1 signal was quantified by sequential masking to DAPI and Iba1. (I) Quantification of STAT1 signal intensity shown in panel H (n = 4 mice per group; *p < 0.05). Data were analyzed using unpaired two-tailed t-tests and are presented as mean ± SEM. Scale bars, 20 µm.

Despite these shared effects on myelin-associated microglial programs, ApoE and Trem2 deficiency differentially impacted microglial lipid handling. While WD feeding increased overall LD abundance within the ME of both *Apoe*⁻^/^⁻ and *Trem*2⁻^/^⁻ mice relative to controls (**Figure 4E-F**), the distribution of LDs within microglia diverged markedly between genotypes. Microglia in the ME of *Apoe*⁻^/^⁻ mice accumulated excess intracellular LDs, whereas microglia in *Trem2*⁻^/^⁻ mice exhibited a near-complete absence of LDs, despite elevated lipid content in surrounding cells (**Figure 4G**). These findings indicate that ApoE and Trem2 reciprocally regulate microglial lipid storage in ME microglia of mice fed a WD. We next examined whether these differences in lipid handling were associated with altered interferon signaling. Remarkably, we found that WD-induced STAT1 induction in ME microglia was preserved in *Apoe*⁻^/^⁻ mice but was markedly attenuated in *Trem*2⁻^/^⁻ mice under WD feeding (**Figures 4H-I**). Thus, although both ApoE and Trem2 contribute to microglial engagement with myelin, Trem2 is selectively required to link lipid-associated cues with interferon-responsive microglial programs in the ME.

Consistent with prior reports^42^, *Apoe*⁻^/^⁻ mice fed a WD gained less body weight than wild-type controls under WD feeding, whereas *Trem2*⁻^/^⁻ mice gained more weight than their respective controls (**Figures S7A–B**). Importantly, genotypes exhibiting greater weight gain did not show greater microglial lipid accumulation or interferon-associated signaling, indicating that these microglial phenotypes are not explained solely by differences in body weight. Given the use of global knockout models, genotype-dependent effects on weight gain cannot be directly attributed to microglial-specific mechanisms.

Together, these data reveal that although *Apoe* and *Trem2* are both required for microglial interactions with myelin in the ME, Trem2 uniquely integrates lipid accumulation with interferon-responsive signaling during dietary challenge.

### Human APOE4 isoform exacerbates microglial lipid overload and impairs myelin organization in the ME during WD feeding

To translate our findings from global *Apoe* loss-of-function models into a more clinically relevant human-centered context, we examined humanized *APOE* knock-in mice expressing either *APOE3/E3* or *APOE4/4* under endogenous regulatory control, with *APOE3/E3* mice serving as a reference genotype and *APOE4/E4* mice reflecting a common human risk allele associated with metabolic and neurodegenerative disease. This approach preserves normal APOE expression levels while assessing the impact of human allelic variation, unlike complete gene deletion.

Microglia in the ME of *APOE4/E4* mice fed a WD exhibited marked increased Clec7a expression compared with *APOE3/E3* controls (**Figures 5A–B**), consistent with increased activation of the myelin responsiveness, and STAT1 immunoreactivity (**Figures 5C–D**), indicating amplified interferon. We next examined the impact of APOE isoform identity on microglial lipid handling in the ME. Compared with *APOE3/E3* controls, microglia from *APOE4/E4* mice fed a WD had greater LD content, reflected by both higher LD volume and higher frequency of LD-laden microglia (**Figures 5E–F**). These findings indicate that APOE isoform identity influences microglial lipid homeostasis in the ME during dietary stress.

**Figure 5.**
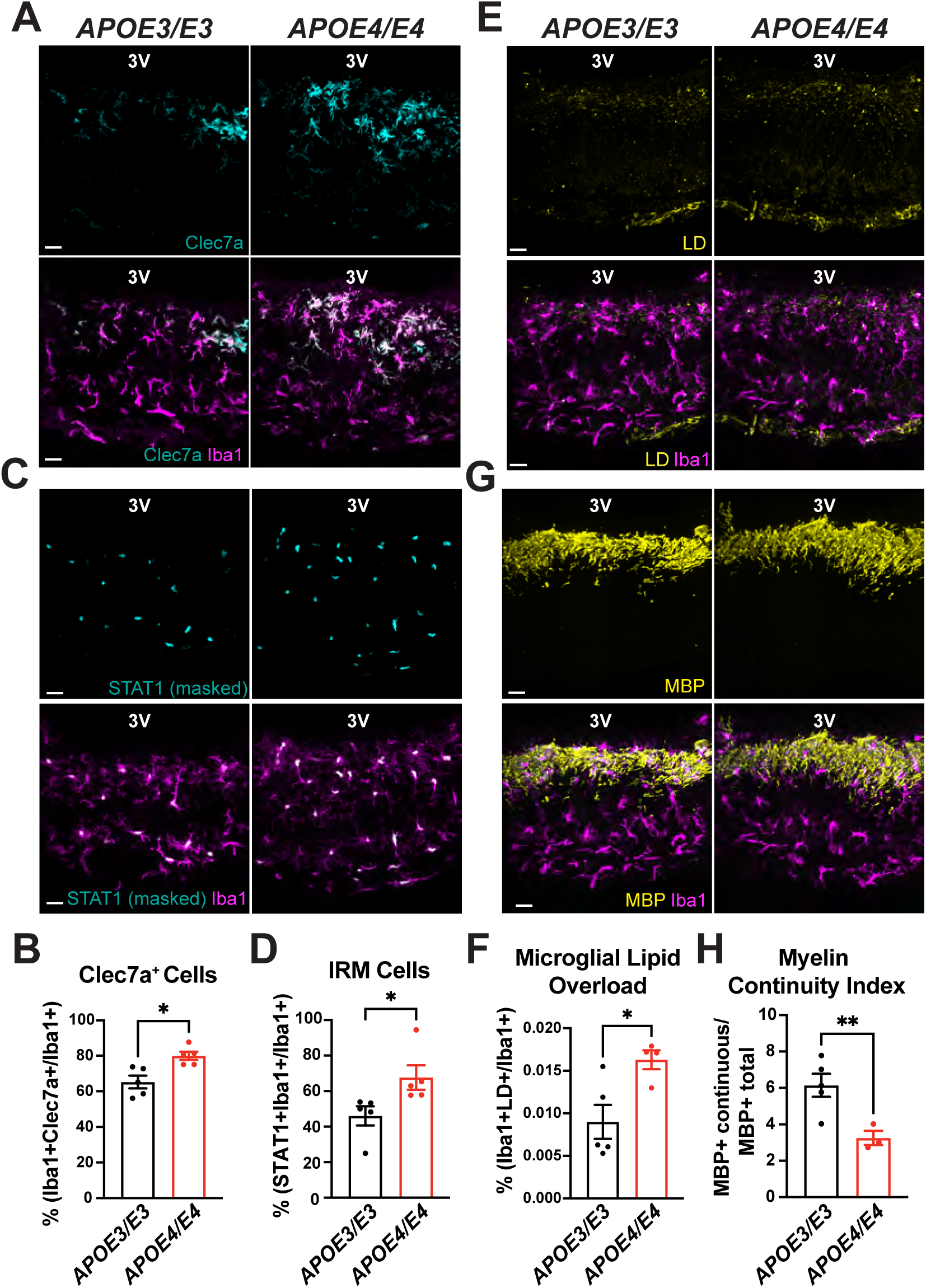
APOE4 enhances interferon signaling, alters microglial lipid handling, and impairs myelin organization in median eminence microglia during western diet feeding. (A) Representative images showing Clec7a expression in Iba1⁺ microglia within the median eminence of adult male humanized APOE3/E3 and APOE4/E4 mice fed a western diet (WD) for 4 weeks. (B) Quantification of microglial Clec7a expression shown in panel A (n = 5 mice per group; *p < 0.05). (C) Representative images showing STAT1 immunoreactivity within Iba1⁺ microglia in the ME of humanized APOE3/E3 and APOE4/E4 mice fed a WD for 4 weeks. STAT1 signal was quantified by sequential masking to DAPI and Iba1. (D) Quantification of STAT1 signal intensity shown in panel C (n = 5 mice per group; *p < 0.05). (E) Representative images of LipidSpot staining used to label lipid droplets (LDs) and myelin basic protein (MBP) staining in Iba1⁺ microglia within the ME of APOE3/E3 and APOE4/E4 mice fed a WD for 4 weeks. (F) Quantification of microglial LD accumulation from the dataset shown in panel E (n = 4–5 mice per group; *p < 0.05). (G) Quantification of myelin continuity from the dataset shown in panel E (n = 3–5 mice per group; **p < 0.01). Data were analyzed using unpaired two-tailed t-tests and are presented as mean ± SEM. Scale bars, 20 µm.

We next assessed myelin organization in the ME. Myelin in the ME of *APOE4/E4* mice fed a WD exhibited reduced structural continuity compared with *APOE3/E3* controls, coinciding with increased microglial LD burden and interferon signaling under dietary stress (**Figures 5G-H**). Consistent with prior reports in male APOE targeted-replacement^43^, *APOE4/E4* mice gained more body weight than *APOE3/E3* controls under WD feeding (**Figure S7C**). Because these models preserve physiological APOE expression and differ only in isoform identity, the observed differences in body weight likely reflect isoform-dependent effects acting across multiple tissues.

### Microglial APOE is required for interferon-associated microglial programs but not for lipid accumulation

To assess the cell-intrinsic role of microglial *APOE*, we used a humanized *APOE3* targeted-replacement mouse line, in which the mouse *Apoe* coding sequence is replaced by the human *APOE3* gene expressed from the endogenous mouse *Apoe* locus and flanked by LoxP sites, enabling conditional deletion upon Cre recombinase expression. These mice were crossed with inducible Cx3cr1-Cre^ER^ mice to generate MG-Apoe^KO^ mice. Cre-mediated recombination resulted in a near-complete loss of *APOE* transcripts in ME microglia (**Figures 6A–B**). Despite this lack of microglial *APOE* transcription, APOE protein remained surprisingly detectable among microglia in the ME of MG-Apoe^KO^, consistent with potential microglial uptake of APOE from non-microglial sources that continue to express *APOE3* within the local brain environment (**Figure 6C**). Thus, this model selectively removes microglia-derived APOE while preserving its availability to microglia via uptake from the extracellular milieu in the ME.

**Figure 6.**
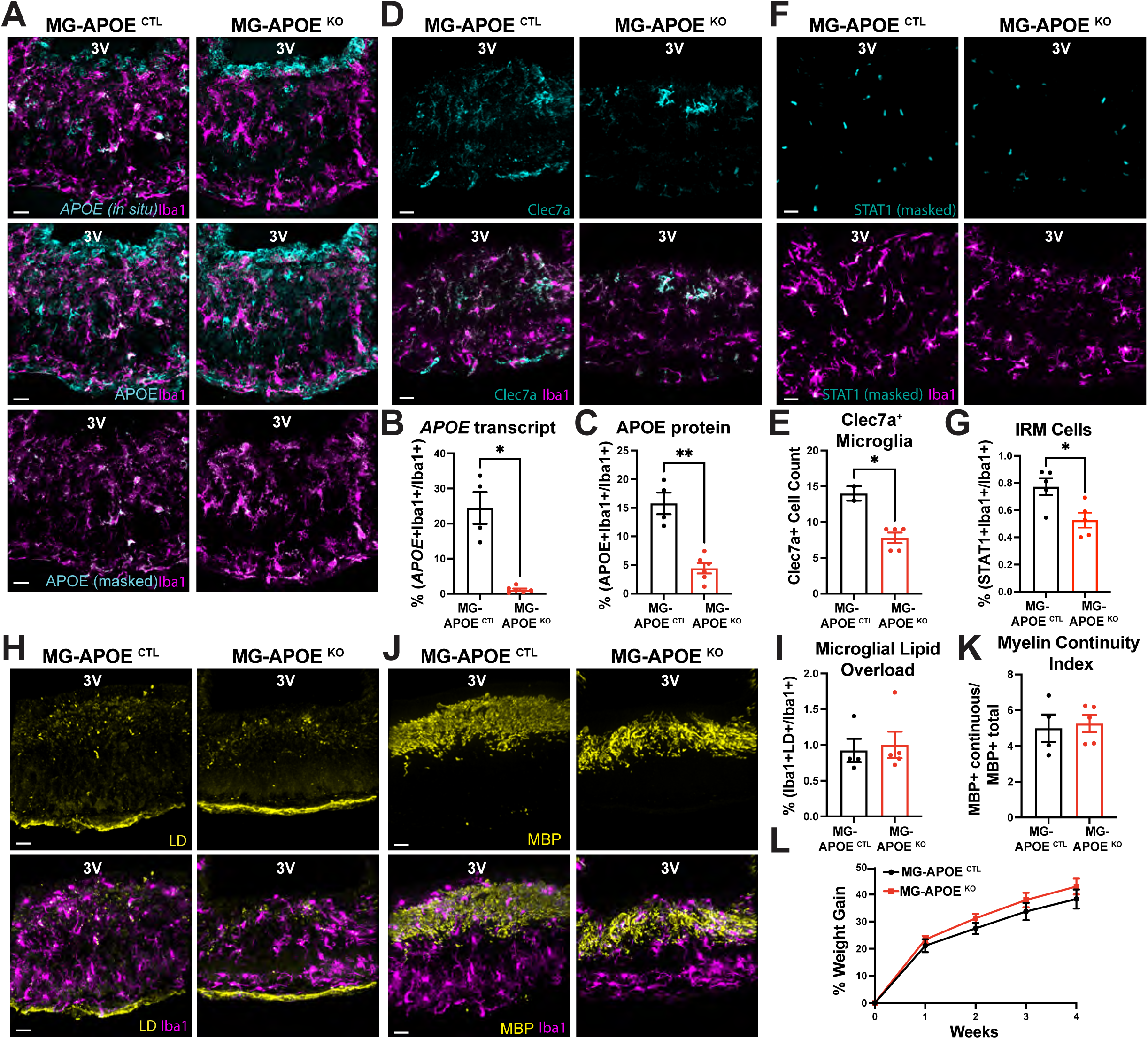
Microglia-derived APOE is required for interferon signaling but dispensable for lipid accumulation and myelin fragmentation in the median eminence. (A) Representative images showing APOE transcripts colocalized with the microglial marker Iba1, APOE protein colocalized with Iba1, and APOE protein signal masked to Iba1 immunostaining in the hypothalamic median eminence of adult male control (MG-APOE ^CTL^) and microglia-specific APOE knockout (MG-APOE ^KO^) mice fed a western diet (WD) for 4 weeks. (B, C) Quantification of APOE transcript and APOE protein signal shown in panel A (n = 4–6 mice per group; *p < 0.05, **p < 0.01). (D) Representative images showing Clec7a expression in Iba1⁺ microglia within the ME of MG-APOE ^CTL^ and MG-APOE ^KO^ mice fed a WD for 4 weeks. (E) Quantification of microglial Clec7a expression shown in panel D (n = 2–5 mice per group; *p < 0.05). (F) Representative images showing STAT1 immunoreactivity within Iba1⁺ microglia in the ME of MG-APOE ^CTL^ and MG-APOE ^KO^ mice. STAT1 signal was quantified by sequential masking to DAPI and Iba1. (G) Quantification of STAT1 signal intensity shown in panel F (n = 5 mice per group; *p < 0.05). (H) Representative images of LipidSpot staining used to label lipid droplets (LDs) in Iba1⁺ microglia within the ME of MG-APOE ^CTL^ and MG-APOE ^KO^ mice. (I) Quantification of microglial LD accumulation shown in panel H (n = 4–5 mice per group). (J) Representative images showing myelin basic protein (MBP) within Iba1⁺ microglia in the ME of MG-APOE ^CTL^ and MG-APOE ^KO^ mice. (K) Quantification of myelin continuity index is shown in panel J (n = 4–5 mice per group). (L) Body weight curves for MG-APOE CTL and MG-APOE KO mice over 4 weeks of WD feeding (n = 4–6 mice per group). Data were analyzed using unpaired two-tailed t-tests and are presented as mean ± SEM. Scale bars, 20 µm.

Microglia in the ME of MG-Apoe^KO^ mice fed a WD had significantly reduced Clec7a expression (**Figure 6D-E**) and markedly attenuated STAT1 induction (**Figures 6F-G**), indicating impaired engagement of both myelin-responsive and interferon-associated microglial programs. In contrast, microglial LD accumulation was unaffected by microglial *APOE* deletion, with no differences seen in either total LD volume or the proportion of lipid-laden microglia when compared with control mice (**Figures 6H–I**). Similarly, myelin organization within the ME was preserved, as reflected by comparable myelin continuity indices between MG-*APOE*^KO^ and MG-*APOE*^CTRL^ mice (**Figure 6J–K**). These findings indicate that microglial *APOE* is required to link myelin-associated cues to interferon-responsive microglial programs in the ME, while LD accumulation itself can occur independently of microglial *APOE* expression. MG-Apoe^KO^ mice gained similar weight to control mice when fed a WD (**Figure 6L**), in the context of preserved microglial LD accumulation and myelin organization.

### Pharmacological restoration of microglial lipid homeostasis ameliorates WD-induced metabolic dysfunction

To determine whether microglial lipid accumulation and associated stress programs are therapeutically reversible, we employed a synthetic HDL-based liver X receptor agonist formulation (sHDL-LXRa) designed for preferential uptake by phagocytic cells. Following systemic administration, fluorescently labeled sHDL particles were robustly taken up by microglia in the ME, with comparatively lower signal detected in neighboring non-microglial cell types (**Figure S8A-B**). Consistent with prior reports using this formulation^44^, sHDL-LXRa treatment did not induce hepatic expression of canonical LXR target genes (**Figure S8C**), confirming that preferential myeloid targeting was achieved without detectable off-target hepatic lipogenic responses.

To test therapeutic efficacy after disease onset, mice were fed a WD for six weeks prior to treatment initiation, a time point sufficient to induce microglial lipid accumulation and interferon-associated programs within the ME. sHDL-LXRa was then administered systemically every other day, allowing assessment of whether targeted restoration of microglial lipid homeostasis can reverse diet-induced hypothalamic dysfunction after it has already manifested. sHDL–LXRa treatment significantly increased the expression of LXR target genes, including *Apoe* and *Abca1*, in ME microglia (**Figure 7C-D**; **Figure S8D-E**), confirming effective activation of microglial LXR signaling in vivo. This activation was accompanied by a marked normalization of microglial lipid handling, evidenced by a robust reduction in LD accumulation within Iba1^+^ ME microglia despite ongoing WD consumption (**Figure 7E–F**). These findings demonstrate that diet-induced microglial lipid stress in the ME is actively reversible by targeting LXR activation. This improvement in lipid handling was accompanied by restored myelin handling and enhanced myelin organization within the ME, as reflected by an increased MBP continuity index (**Figure 7G–I**). In parallel, engagement of stress-associated microglial programs was attenuated, with marked reductions in Clec7a and STAT1 expression in ME microglia (**Figure 7J–M**). Thus, although LD accumulation can persist in the absence of APOE-dependent interferon and myelin-responsive programs, therapeutic restoration of microglial lipid homeostasis coordinately attenuates both responses, supporting a model in which microglial-intrinsic *APOE* couples lipid stress to coordinated transcriptional states downstream of LXR activation.

**Figure 7.**
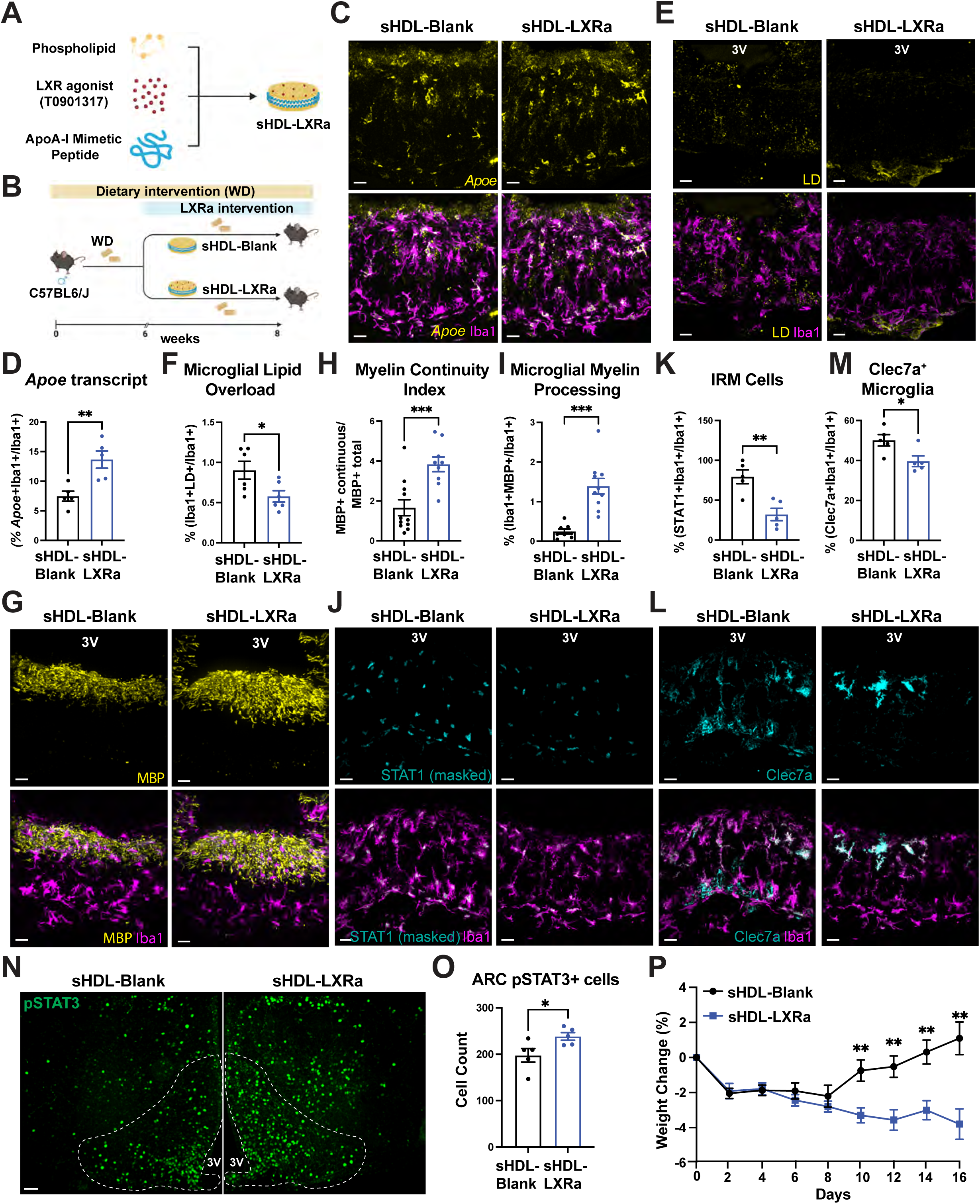
Targeted sHDL–LXR agonist restores microglial lipid handling in the median eminence, improves leptin responsiveness, and attenuates western diet–induced body-weight gain. (A) Schematic of the synthetic sHDL–LXRa formulation. (B) Schematic of the in vivo delivery paradigm for sHDL–LXRa or vehicle. (C) Representative images showing Apoe transcripts colocalized with the microglial marker Iba1 in the hypothalamic median eminence (ME) of adult male mice treated with vehicle or sHDL–LXRa. (D) Quantification of microglial *Apoe* expression shown in panel C (n = 5 mice per group; **p < 0.01). (E) Representative images of LipidSpot staining used to label lipid droplets (LDs) in Iba1⁺ microglia within the ME of vehicle- or sHDL–LXRa–treated mice. (F) Quantification of microglial LD accumulation shown in panel E (n = 6 mice per group; *p < 0.05). (G) Representative images showing myelin basic protein (MBP) within Iba1⁺ microglia in the ME of vehicle- or sHDL–LXRa–treated mice. (H,I) Quantification of myelin continuity index and microglial myelin processing is shown in panel G (n = 10 mice per group; ***p < 0.001). (J) Representative images showing STAT1 immunoreactivity within Iba1⁺ microglia in the ME of vehicle- or sHDL–LXRa–treated mice. STAT1 signal was quantified by sequential masking to DAPI and Iba1. (K) Quantification of STAT1 signal intensity shown in panel J (n = 5 mice per group; **p < 0.01). (L) Representative images showing Clec7a expression in Iba1⁺ microglia within the ME of vehicle- or sHDL–LXRa–treated mice. (M) Quantification of microglial Clec7a expression shown in panel L (n = 5 mice per group; *p < 0.05). (N) Representative images showing phosphorylated STAT3 (pSTAT3) in the hypothalamus of vehicle- or sHDL–LXRa–treated mice following leptin administration. (O) Quantification of leptin-induced pSTAT3 signal shown in panel N (n = 5 mice per group; *p < 0.05). (P) Body weight curves for chronically western diet–fed mice treated with sHDL–LXRa or vehicle from treatment initiation to tissue collection (n = 10 mice per group). Data were analyzed using unpaired two-tailed t-tests and are presented as mean ± SEM. Scale bars, 20 µm, except panel N (50 µm). 3V, third ventricle.

We next examined whether LXR-mediated microglial metabolic reprogramming within the ME translated into functional improvements in hypothalamic signaling. sHDL–LXRa–treated mice displayed enhanced leptin responsiveness, as indicated by an increased number of pSTAT3⁺ neurons following leptin administration compared to vehicle-treated WD controls (**Figure 7N–O**). Notably, these central effects were accompanied by a significant attenuation of WD-induced body weight gain over the treatment period (**Figure 7P**), indicating that modulation of microglial lipid metabolism is associated with improved energy homeostasis even after disease establishment.

Collectively, these data establish microglial lipid dysregulation at the hypothalamic neurovascular interface as a mechanistically relevant consequence of WD exposure that contributes to hypothalamic dysfunction. Importantly, this state is reversible and therapeutically actionable. Targeted restoration of microglial lipid homeostasis suppresses stress-associated microglial programs, improves myelin organization, restores hypothalamic leptin responsiveness, and limits diet-induced weight gain. Together, these findings position microglial lipid metabolism within the ME as a central control node through which nutritional stress reshapes hypothalamic signaling and systemic metabolic homeostasis.

## Discussion

In this study, we define microglial lipid metabolism as a central and dynamically regulated determinant of hypothalamic function during dietary excess. Focusing on the ME, a specialized neurovascular interface within the MBH, we show that WD feeding engages coordinated microglial programs characterized by intracellular lipid accumulation together with Clec7a- and interferon-associated signaling. Importantly, our data indicate that ME myelin disruption is not a uniform or inevitable consequence of WD exposure. Instead, chronic dietary excess establishes a permissive environment for myelin vulnerability, with disruption emerging when microglial lipid handling capacity is exceeded, and stress-associated signaling becomes dominant. Together, these findings position hypothalamic microglia as active regulators of lipid processing, tissue vulnerability, and metabolic signaling rather than passive responders to nutritional stress.

A key conceptual advance of this work is a shift from descriptive notions of microglial “reactivity” toward the delineation of distinct transcriptional programs allowing for the construction of a functionally interpretable state-based cellular framework. Microglial responses in metabolic disease have often been inferred from generic markers or morphological changes, leaving unresolved what these cells are functionally engaged in during nutritional stress. Here, we show that WD feeding does not elicit a unitary inflammatory response, but instead drives distinct microglial programs linked to lipid handling, myelin engagement, and interferon signaling. Notably, these programs are not strictly coupled: microglial lipid accumulation and changes in myelin organization can occur in the absence of interferon-associated transcriptional activation, supporting the view that stress-associated signaling reflects a regulated microglial state transition rather than a passive consequence of lipid exposure. This dissociation is particularly evident in our analysis of microglial ApoE function. Loss of cell-intrinsic *APOE* expression did not eliminate APOE protein from microglia, consistent with prior observations that microglia can acquire APOE protein from extrinsic sources^45^. Despite preserved APOE protein availability and ongoing LD accumulation, microglial *APOE* deletion abolished Clec7a induction and interferon-associated signaling. These findings demonstrate that lipid exposure and myelin engagement alone are insufficient to drive stress-associated microglial programs in the absence of intrinsic APOE expression and suggest that interferon-associated responses may integrate lipid stress signals beyond myelin-derived substrates. Instead, these findings are consistent with a model in which extrinsic APOE can contribute to microglial lipid handling, whereas microglia-intrinsic APOE is regulated for engagement of stress-associated transcriptional programs, helping to explain why lipid accumulation and interferon signaling do not necessarily scale together.

Our study further reveals pronounced anatomical specificity in hypothalamic microglial responses to dietary stress. Although the MBH is often treated as a uniform metabolic hub, microglia in the ME exhibit distinct lipid-handling and interferon-associated programs that are largely absent from adjacent hypothalamic regions. This specialization likely reflects the unique vascular and extracellular environment of the ME, which exposes resident microglia to circulating lipids and hormones, and inflammatory cues, while necessitating continuous structural remodeling. Rather than being uniformly disrupted by a chronic WD feeding, myelin integrity within the ME appears sensitive to the relative balance of microglial programs, becoming vulnerable when lipid-handling capacity is exceeded and stress-associated programs become dominant. These findings identify the ME as a discrete microglial niche within the MBH and underscore the importance of subregional resolution when interpreting hypothalamic microglial responses in metabolic disease models.

Within this niche, we observe evidence of a constitutive interferon signaling landscape. Even under baseline conditions, STAT1 immunoreactivity is readily detectable within the ME, suggesting that a basal interferon tone is a feature of the local ME environment rather than a response uniquely induced by dietary stress. Given the strategic positioning of the ME at the blood–CSF interface and its roles in nutrient sensing, transport, and barrier plasticity, tonic interferon signaling cues within this niche may shape this immune–metabolic context encountered by resident microglia. In this framework, WD feeding does not initiate interferon signaling *de novo* but instead alters how microglia engage a pre-existing interferon environment, promoting integration of lipid stress from multiple sources with microglial transcriptional programs. This framework provides a potential explanation for both the anatomical restriction of interferon-associated microglial states to the ME and their sensitivity to genetic context, including microglia-intrinsic APOE function.

An additional implication of these findings is that ME microglia may indirectly influence hypothalamic circuits by regulating how lipid and inflammatory cues are processed within the ME microenvironment. Given that the ME serves as a gateway through which peripheral signals reach neurons in the ARC, alterations in microglial lipid metabolism and stress-associated programs within the ME can reshape the extracellular and barrier properties through which circulating metabolic signals access ARC circuits. In this framework, ME microglia do not need to directly engage neuronal synapses to influence metabolic control; instead, their capacity to buffer lipid and inflammatory stress within the ME may indirectly modulate ARC function by altering the quality and fidelity of peripheral signal transmission. This interpretation is consistent with our observation that reprogramming ME microglia improves hypothalamic leptin responsiveness without requiring direct microglial modulation of neuronal identity or activity.

Beyond these barrier-associated mechanisms, our data also highlight an underappreciated role for microglia-oligodendrocyte interactions in shaping ME vulnerability under dietary stress. Unlike classical white matter tracts, the ME contains sparse but dynamically remodeled myelin that is continuously exposed to metabolic and inflammatory cues. Microglia play a central role in the clearance, recycling, and redistribution of myelin-derived lipids, thereby positioning them as key regulators of oligodendrocyte function and myelin maintenance.

In this context, WD-induced lipid accumulation within microglia is not merely a marker of stress but likely reflects altered handling of myelin-associated lipid substrates within a broader lipid-rich microenvironment. These observations indicate that myelin disruption within the ME is not a passive consequence of dietary excess or inflammatory tone alone but instead reflects context-specific microglial lipid-myelin interactions.

In this context, ApoE and Trem2 emerge as key but non-redundant regulators of microglial state. While both pathways have been extensively studied in neurodegenerative contexts, our data extend their relevance to metabolic stress in the hypothalamus. Trem2 primarily governs lipid uptake and sensing, whereas ApoE influences lipid handling and stabilization of stress-associated transcriptional programs. Expression of the human *APOE4* isoform amplified WD-induced microglial lipid dysregulation, interferon signaling, and myelin vulnerability, indicating that *APOE4* biases the magnitude and persistence of microglial responses under metabolic challenge. These findings support a model in which ApoE and Trem2 function together to shape microglial adaptation to lipid-rich environments, while in humans, *APOE* genetic variation may tune the balance between adaptive and maladaptive outcomes.

Our results further demonstrate that rather than being fixed, WD-induced microglial states are pharmacologically tractable. Targeted delivery of an sHDL–LXR agonist after prolonged WD feeding restored microglial lipid homeostasis, improved myelin organization, and attenuated interferon-associated microglial programs within the ME. These cellular effects were accompanied by enhanced hypothalamic leptin responsiveness and reduced weight gain, indicating that microglial reprogramming within the ME can translate into functional improvements in metabolic signaling. The absence of canonical hepatic lipogenic responses to sHDL-LXRa treatment further supports the specificity of this approach, distinguishing it from systemic LXR activation. Notably, the absence of a body weight phenotype after microglia-specific *APOE* deletion contrasts with the metabolic improvement observed after sHDL-LXRa treatment, indicating that broad restoration of microglial lipid homeostasis across diverse lipid inputs, rather than modulation of microglial APOE alone, is required to influence systemic metabolic outcomes.

An intriguing implication of these findings is a potential reciprocal relationship between interferon signaling and LXR-dependent lipid metabolism in ME microglia. Prior work in myeloid and brain-resident cells has shown that STAT1 activation can antagonize LXR-dependent transcription, while LXR activation suppresses interferon-responsive transcriptional programs^46,47^, suggesting that interferon-associated microglial states actively reprogram lipid handling rather than simply reflecting lipid stress. This relationship mirrors antiviral immune responses, in which interferon signaling drives redistribution of accessible cholesterol to restrict pathogen replication^48,49^. By analogy, sustained interferon engagement in ME microglia may redirect lipid handling under dietary stress, with downstream consequences for myelin-associated lipid processing and hypothalamic function. Future studies are warranted to directly test whether interferon signaling is sufficient and/or necessary to drive lipid dysregulation in ME microglia.

Finally, our findings may inform understanding of interindividual variability in metabolic vulnerability across APOE genotypes. Although *APOE4* is best known for its association with neurodegenerative disease, emerging human studies report context-dependent associations between *APOE* genotype and metabolic traits, including lipid profiles, insulin sensitivity and body weight^50–52^. In this context, *APOE* genotype may shape how the central nervous system adapts to lipid-rich or inflammatory environments. Together, these findings provide a cellular framework through which *APOE* genetic variation could influence metabolic resilience across nutritional and inflammatory contexts.

Together, these findings raise important questions regarding how microglial lipid metabolism shapes hypothalamic function under metabolic stress. Defining how diet-responsive microglial state transitions within the ME influence barrier properties, glial–glial interactions, and access of circulating metabolic signals to hypothalamic circuits will be critical for understanding how diet alters central sensing of peripheral cues. Moreover, determining the temporal stability of these microglial programs and whether early dietary exposures imprint long-lasting changes in ME microglia may provide insight into the developmental programming of metabolic disease. By defining microglial lipid metabolism as a dynamic regulator of ME vulnerability and metabolic signaling, this work positions microglia as active participants in diet-induced hypothalamic dysfunction and highlights new opportunities for targeted modulation of brain–metabolic interfaces.

### Limitations

Several limitations of this study warrant consideration. First, although we used an inducible Cx3cr1-Cre^ER^ strategy to delete *APOE* in microglia, this approach does not exclusively target microglia, as Cx3cr1 is also expressed by non-microglial myeloid populations, including border-associated macrophages. While a washout period was designed to minimize recombination in short-lived cells, contributions from long-lived macrophage populations cannot be fully excluded. Notably, microglia can acquire APOE from non-microglial sources, meaning that phenotypes driven by APOE availability may be masked *when APOE* is deleted in only a single cell compartment. Accordingly, although our findings strongly support a role for microglia-intrinsic APOE in regulating interferon-associated programs within the ME, definitive attribution to microglia alone will require complementary approaches that disentangle microglial *APOE* expression from extracellular APOE availability.

Second, although the sHDL–LXR agonist was designed to preferentially target phagocytic cells and we provide evidence of selective uptake by ME microglia, the compound was delivered systemically, and off-target effects in peripheral tissues cannot be entirely excluded. Importantly, we did not observe induction of canonical hepatic lipogenic gene programs typically associated with systemic LXR activation, supporting a degree of central and cellular specificity. Nevertheless, future studies employing brain-restricted or microglia-specific delivery strategies will be necessary to further refine cell-autonomy and distinguish between direct and indirect contributions to the observed metabolic improvements.

Third, our analyses focused primarily on Western diet feeding and the ME, and it remains to be determined whether similar microglial state transitions occur under other metabolic challenges or within additional hypothalamic niches. Finally, extending these findings across sex, age, and developmental windows will be important for defining the broader physiological relevance of microglial lipid metabolism in metabolic disease.

## Acknowledgements

This work was supported by Larry L. Hillblom Foundation (Start-Up grant to M.V.), the NIDDK (R01 DK134782-01 to M.V. and R01 DK103175 to S.K). Metabolic phenotyping was supported by the Nutrition Obesity Research Center at UCSF (DK098722). A spinning confocal microscope was purchased with an NIH S10 grant (S10OD028611). We thank CureAlz for generously providing the humanized *APOE3* and *APOE4* targeted-replacement mice, and the UCSF Flow Cytometry Core for technical support.

## Author Contributions

E.R.M. designed and performed most experiments, analyzed data, contributed to the conceptual development of the study, and edited the manuscript. A.F. performed selected experiments and contributed to conceptual discussions. L.M. formulated and characterized the synthetic HDL nanoparticles. S.M.B.M., V.P., and S.P. provided technical assistance with experiments. M.K.C. assisted with single-cell RNA-sequencing analyses. R.T.C. assisted with experimental work. R.W.B. performed fluorescence-activated cell sorting. A.S. provided nanoparticle reagents and technical expertise. V.N. performed single-cell RNA-sequencing analysis and generated associated figures. S.K.K. provided funding support, analyzed data, and edited the manuscript. M.V. conceived and supervised the study, led the conceptual development of the work, provided funding support, analyzed data, and both wrote and edited the original manuscript.

## STAR METHODS

## RESOURCE AVAILABILITY

### Lead contact

Further information and requests for resources and reagents should be directed to and will be fulfilled by the Lead Contact, Martin Valdearcos.

### Material availability

This study did not generate new unique reagents.

### Data and code availability

Single-cell RNA-sequencing data generated in this study will be deposited in a public repository and will be made available upon publication.

## EXPERIMENTAL MODEL AND STUDY PARTICIPANT DETAILS

### Animal Husbandry

All mouse strains were maintained in the University of California, San Francisco (UCSF) specific pathogen–free animal facility. All animal procedures were performed in accordance with the NIH Guidelines for the Care and Use of Laboratory Animals and were approved by the UCSF Institutional Animal Care and Use Committee and the Laboratory Animal Resource Center. Mice were housed under a 12-hour light/dark cycle (lights on from 07:00 to 19:00) at 20–26 °C and 30–70% humidity. Animals were group-housed when possible (up to five mice per cage) with ad libitum access to water. Littermate controls were used whenever feasible for all experiments.

### Rodent Lines and Experimental Animals

The following mouse strains were used: C57BL/6J (The Jackson Laboratory, #000664); B6.129P2-Apoe^tm1Unc/J^ (The Jackson Laboratory, #002052); C57BL/6J-Trem2^em2Adiuj^/J (The Jackson Laboratory, #027197); B6.Cg-Apoe^em2(APOE*)Adiuj/J^ (The Jackson Laboratory, #029018); C57BL/6J-Apoe^em8(APOE*)Msasn/J^ (The Jackson Laboratory, #038051); B6.129P2(C)-Cx3cr1^tm2.1(cre/ERT2)Litt^/ WganJ (The Jackson Laboratory, #021160); Humanized APOE3 targeted-replacement mice, in which the coding sequence of the endogenous murine *Apoe* gene is replaced by the human APOE3 sequence flanked by loxP sites to enable conditional disruption of *APOE* expression (provided through the Cure Alzheimer’s Fund) and PDGFRα-CreER::Myrf fl/fl mice obtained from the laboratory of Dr. Jonah Chan. Anesthesia was performed using isoflurane or Avertin for terminal procedures.

### Generation of Experimental Mice

To achieve microglial deletion of human *APOE* gene, heterozygous CX3CR1^CreER/+^ mice were crossed with *APOE3*^fl/fl^ generating CX3CR1^CreER/+^::APOE3^fl/fl^ offspring. These mice are hereafter referred to as MG-*APOE*^KO^. Cre-negative APOE3^fl/fl^ littermates, which retain intact human *APOE* expression, served as controls and are hereafter referred to as MG-*APOE*^CTL^. For oligodendrocyte-lineage-specific deletion of *Myrf,* heterozygous PDGFRα^CreER/+^ mice were crossed with Myrf ^fl/fl^ to generate PDGFRα ^CreER/+ fl/fl^ mice are referred to as OPC-Myrf^KO^, with Cre-negative Myrf ^fl/fl^ littermates, which are referred to as OPC-Myrf^CTL^, serving as controls. C57BL/6J mice were used as controls for whole-body knockout experiments. All experimental mice were 8-20 weeks of age at the time of analysis.

### Induction of CreER-Mediated Recombination

To induce CreER-mediated genetic recombination, 20 mg/mL tamoxifen (MP Biomedicals) dissolved in corn oil (Sigma-Aldrich) was delivered via enteric gavage to all experimental and control mice at 8w of age for 3 (Cx3cr1^CreER/+::^*APOE*3^fl/fl^) or 5 (PDGFRα^CreER/+^::Myrf^fl/fl^) consecutive days. For studies using PDGFRa^CreER/+^::Myrf^fl/fl^ mice, mice were maintained on chow diet for an additional week to allow for full genetic recombination prior to dietary challenge. For studies using Cx3cr1^CreER/+^::*APOE*3^fl/fl^ mice, adult mice received tamoxifen via enteric gavage, maintained on low-fat diet control for 4 weeks to allow peripheral monocyte turnover prior to diet challenge.

### Dietary Interventions

Adult mice were maintained on a low-fat diet control (CD; LabDiet 5053) or challenged with ad libitum western diet (42% kcal fat, TD. 88137; Envigo) when indicated. For LXR agonism experiments, mice were maintained on a Western diet for 6 weeks prior to the start of the LXR paradigm and continued to have access to ad libitum Western diet throughout the treatment.

## METHOD DETAILS

### Single-cell dissociation and FACS purification

Mediobasal hypothalamic (MBH) tissue was rapidly dissected from adult mice on ice and immediately transferred to ice-cold Hibernate-A medium. Tissue was mechanically dissociated at 4°C using manual Dounce homogenization (10 strokes with a loose pestle, followed by 10 strokes with a tight pestle) to maximize cell recovery while preserving viability. Dissociation buffers were supplemented with RNase inhibitors (SUPERase-In, Invitrogen) and actinomycin D (3 µg/mL) to minimize ex vivo transcriptional activation. All steps were performed on ice or at 4 °C to preserve RNA integrity^1^. Single-cell suspensions were passed through a 40 µm cell strainer and stained with fluorophore-conjugated antibodies against CD11b, along with DAPI to exclude non-viable cells. Live (DAPI⁻), CD11b⁺ cells were isolated by fluorescence-activated cell sorting (FACS). For each biological replicate, exactly 2,000 viable CD11b⁺ cells were collected per mouse to ensure equal input across samples. For single-cell RNA sequencing, MBH tissue was collected from 5 mice fed a Western diet and 5 mice fed a control low-fat diet.

### Single-Cell Library Preparation, Sequencing and Computational Analysis

Sorted cells were labeled with lipid-modified DNA barcodes according to the MULTI-seq protocol to enable sample multiplexing prior to library preparation^33^. Barcoded cells were processed using the 10x Genomics Chromium 3′ v3 chemistry and sequenced on an Illumina NovaSeq 6000 platform to a mean depth of approximately 50,000 reads per cell. Sequencing data were demultiplexed using the kITE pipeline (kallisto indexing and tag extraction)^53,54^ and aligned to the mm10 mouse reference genome using kallisto bustools^55,56^. Raw count matrices were processed and analyzed using Scanpy^57^. High-quality cells were retained based on the following criteria: (1) total UMI counts > 2,300, (2) > 1,150 genes detected, and (3) mitochondrial gene content < 5%. Thresholds were determined by manual inspection of quality control metrics. After filtering, the dataset comprised 6,834 cells. Expression values were log-normalized to 10,000 counts per cell (CP10K), and highly variable genes (n = 5,000) were selected. Dimensionality reduction was performed using principal component analysis, and the resulting low-dimensional representations were used to construct a k-nearest neighbors’ graph (k = 15). Cells were clustered using the Leiden community detection algorithm (resolution = 0.5) and visualized using UMAP, yielding 11 initial clusters.

Two small clusters enriched for neuronal markers and one low-quality cluster dominated by ribosomal gene expression were excluded, resulting in a final dataset of 6,458 myeloid cells for downstream analyses. Differential gene expression analysis was performed using the Wilcoxon rank-sum test to identify cluster-specific markers, and cell types were annotated manually based on canonical gene expression signatures. Four transcriptionally related homeostatic microglial clusters were merged to generate the cell-type annotations shown in Fig. 1B. Differential expression analysis was repeated using these consolidated annotations. The top 10 marker genes for each cluster with log fold change > 1 are shown in Fig. 1C. To further resolve heterogeneity within Apoe⁺ microglia, clustering was repeated at higher resolution (Leiden resolution = 2). Gene signatures corresponding to disease-associated microglia (DAM)^4^, interferon-responsive microglia (IRM)^5^, and the SenMayo senescence program ^58^were used to compute per-cell enrichment scores using the Scanpy score_genes function with default parameters.

### Reagents

The liver X receptor (LXR) agonist T0901317 (T0) was purchased from Cayman Chemical (Ann Arbor, MI). The ApoA-I mimetic peptide 22A (PVLDLFRELLNELLEALKQKLK) was synthesized by GenScript Inc. (Piscataway, NJ). 1,2-dimyristoyl-sn-glycero-3-phosphocholine (DMPC) was purchased from NOF America Corporation (White Plains, NY). Synthetic high-density lipoprotein (sHDL) nanodiscs loaded with the LXR agonist T0 (hereafter referred to as sHDL-LXRa) and corresponding unloaded nanodiscs (sHDL control) were formulated by the laboratory of Dr. Anna Schwendeman (University of Michigan College of Pharmacy). All additional reagents were of analytical grade and obtained from commercial sources.

### Synthesis of sHDL-LXRa and sHDL Control Nanodiscs

sHDL control (vehicle) and sHDL-LXRa nanodiscs were prepared by the Schwendeman laboratory as previously described. For sHDL control nanodiscs, 22A and DMPC were briefly dissolved in acetic acid and combined in glass scintillation vials at a 1:2 (w/w) ratio. For sHDL-LXRa nanodiscs, 22A, DMPC, and T0 were dissolved in acetic acid and combined at a 1:2:0.5 (w/w/w) ratio. Formulations were flash frozen in liquid nitrogen and lyophilized overnight.

Lyophilized powders were rehydrated in 3 mL of phosphate-buffered saline (PBS; pH 7.4), yielding final concentrations of 12 mg/mL 22A, 36 mg/mL DMPC, and 600 µg/mL T0 for sHDL-LXRa formulations. Reconstituted samples were subjected to three heat–cool cycles (45 °C followed by incubation on ice, 5 min each). After the first cycle, formulations were adjusted to pH 6.5–7.0 using 1 N NaOH. Nanodiscs were sterile filtered through 0.22 µm polyethersulfone filters under aseptic conditions. Final formulations were clear and colorless and were stored at 4 °C for up to one week or aliquoted and stored at −20 °C for long-term storage. For fluorescent labeling, DID-labeled sHDL nanodiscs were prepared by incorporating 0.2% (n/n) DID into the DMPC lipid component during nanodisc formulation.

### Characterization of sHDL Nanodiscs

Particle size and monodispersity were assessed by dynamic light scattering (DLS). Encapsulation of T0 within sHDL-LXRa nanodiscs was quantified by ultra-performance liquid chromatography (UPLC). Free T0 was removed using a 7 kDa molecular weight cutoff desalting column prior to analysis. Encapsulation efficiency was calculated by comparing T0 peak areas in desalted versus total samples and ranged from approximately 70–90%.

### LXR Agonist Administration

Mice received intraperitoneal injections of sHDL-LXRa or sHDL control (vehicle) at a dose equivalent to 1.5 mg/kg T0 every other day. Animals were euthanized within 24 hours of the final dose. Brains were collected, fixed in 4% paraformaldehyde, and embedded in M-1 Embedding Matrix for downstream histological analyses.

### Hypothalamic leptin sensitivity

Hypothalamic leptin sensitivity was assessed by measuring leptin-induced STAT3 phosphorylation in the hypothalamus. Mice were fasted for 6 hours (ZT0–ZT6) and subsequently administered an intraperitoneal injection of recombinant mouse leptin (3 mg/kg body weight) or vehicle. Forty-five minutes after injection, mice were anesthetized and transcardially perfused for brain collection. Hypothalamic pSTAT3 immunoreactivity was quantified as described below in the Image Analysis section.

### Microglial histological analysis and confocal microscopy

Mice were anesthetized and transcardially perfused with saline followed by 4% paraformaldehyde (PFA) in phosphate-buffered saline (PBS). Brains were dissected, post-fixed overnight in 4% PFA at 4 °C, and cryoprotected in 30% (w/v) sucrose in PBS. Hypothalamic tissue was embedded in M-1 Embedding Matrix (Epredia, 1310), rapidly frozen on dry ice, and stored at −20 °C until sectioning. Coronal hypothalamic sections (35 µm thick) were cut on a cryostat and processed as free-floating sections. Sections were blocked for 1 h at room temperature in antibody buffer containing 10% normal goat or donkey serum and 0.1% Triton X-100, followed by incubation with primary antibodies diluted in antibody buffer containing 0.1% Triton X-100 overnight at 4 °C. Primary antibodies included Iba1 (1:2,000, guinea pig, Synaptic Systems), STAT1 (1:500, rabbit, Cell Signaling Technology), CD11c (1:500, Armenian hamster, Bio-Rad), myelin basic protein (MBP; 1:1,000, chicken, Aves Labs), Clec7a (1:500, rat, InvivoGen), ApoE (1:500, rabbit, Cell Signaling Technology), phosphorylated STAT3 (pSTAT3; 1:500, rabbit, Cell Signaling Technology), TREM2 (1:500, sheep, R&D Systems), CD68 (1:500, rat, Bio-Rad), GFAP (1:1,000, mouse, Cell Signaling Technology), and Sox10 (1:1,000, goat, R&D Systems). Sections were incubated with appropriate secondary antibodies (1:300; see Key Resources Table) for 1 h at room temperature. For LipidSpot analysis, sections were blocked and incubated with primary antibodies in buffer containing 0.05% Triton X-100, followed by incubation with a secondary antibody cocktail containing LipidSpot 488 (1:1,000) for 2 h at room temperature. Sections were counterstained with DAPI (1:10,000; Thermo Fisher Scientific) and mounted onto glass slides using ProLong Gold antifade reagent. Images were acquired using a Nikon SoRa spinning disk confocal microscope with 20× and 60× objectives (z-step size, 0.5 µm; total z-depth, 25 µm), with laser power and detector gain maintained constant across all samples. One representative image per sample was acquired for analysis

### Z-stack acquisition and STAT1-based microglial analysis

Z-stack images were acquired in the median eminence using a Nikon SoRa spinning disk confocal microscope with a 60× objective (total z-depth, 25 µm; z-step size, 0.5 µm). Image analysis was performed using Imaris software (Oxford Instruments, v10.0.0). DAPI and Iba1 signals were used to render microglia in three dimensions using the Surfaces tool, with rendering parameters maintained consistently across all images. STAT1 signal was sequentially masked first to DAPI and then to microglial 3D surfaces using uniform thresholding parameters. Microglia containing STAT1-masked signal were manually counted and expressed as a fraction of the total microglial population per image. The total microglial number was determined based on cell body counts using unbiased colocalization of DAPI and Iba1 signals, in conjunction with cellular morphology. Border-associated macrophages were manually excluded from all analyses and were defined by Iba1 expression, amoeboid morphology, and localization at the tissue border.

### Myelin Basic Protein volume, continuity index, and microglial engulfment

Images were analyzed using Imaris software (Oxford Instruments, v10.0.0). Three-dimensional surface renderings of microglia and myelin basic protein (MBP) were generated using the Surfaces tool, with segmentation and thresholding parameters maintained consistently across all images. Border-associated macrophages were manually excluded from all analyses as described above. Microglial engulfment of myelin was quantified as the volume of MBP signal overlapping with microglial 3D surfaces, normalized to the total microglial volume per image. Myelin continuity was assessed by calculating a myelin continuity index, defined as the ratio of total MBP 3D surface volume to the number of discrete MBP surfaces per image, with lower values indicating greater myelin fragmentation.

### Immunoreactive Cell Quantification

Images were analyzed using Imaris software (Oxford Instruments, v10.0.0). Total microglial number was determined using the Imaris Spots tool based on unbiased colocalization of DAPI and Iba1 signal. Border-associated macrophages were manually excluded from all analyses. Clec7a and CD11c signals were thresholded using parameters maintained consistently across all images and quantified using the Imaris Spots tool when colocalized with identified microglia. Expression was reported as the proportion of Clec7a⁺ or CD11c⁺ microglia relative to the total microglial population per image. pSTAT3-positive cells were identified using the Imaris Spots tool based on colocalization between the pSTAT3 signal and DAPI, with exposure thresholds maintained consistently across samples. Quantification was restricted to the arcuate nucleus, which was defined in an unbiased manner based on DAPI nuclear architecture.

### Lipid Droplet (LD) Volume and Microglial Lipid Overload

Images were analyzed using Imaris software (Oxford Instruments, v10.0.0). Microglia and lipid droplets (LDs) were identified by three-dimensional surface rendering of Iba1 and LipidSpot signals, respectively, using the Surfaces tool, with segmentation and thresholding parameters maintained consistently across all images. LipidSpot labeling was used as a surrogate marker for neutral lipid droplets. Total LD volume was quantified per image. Microglial lipid overload was calculated as the volume of LD (LipidSpot) signal overlapping with microglial 3D surfaces, normalized to the total microglial surface volume per image. Border-associated macrophages were manually excluded from all analyses as described above. To assess LD accumulation as a function of microglial activation state, the CD11c signal was rendered in three dimensions, with border-associated macrophages excluded based on tissue border localization and amoeboid morphology. Microglia were classified as CD11c⁺ or CD11c⁻ based on a thresholded CD11c surface-to-cell volume ratio, with classification parameters maintained consistently across images. For each image, LD accumulation was quantified as the volume of LD signal overlapping with CD11c⁺ or CD11c⁻ microglial surfaces, normalized to the corresponding total microglial volume. To evaluate LD accumulation in relation to interferon signaling, microglia were manually classified as STAT1⁺ or STAT1⁻ based on STAT1 immunoreactivity as described above. The volume of LD signal overlapping with STAT1⁺ or STAT1⁻ microglial surfaces was quantified and expressed as a proportion of total LD volume engulfed by microglia.

### Imaris-based 3D surface rendering and volume analysis

Images were analyzed using Imaris software (Oxford Instruments, v10.0.0). Trem2, CD68, STAT1, CD11c, and myelin basic protein (MBP) signals were reconstructed by three-dimensional surface rendering using the Surfaces tool, with segmentation and thresholding parameters maintained consistently across all samples. Total Trem2 signal was quantified per image within the ME. For CD68, STAT1, CD11c, and MBP, hypothalamic regions of interest were defined manually in an unbiased manner based on DAPI nuclear architecture and signal volume was quantified within these regions.

### RNAscope Fluorescent In-situ Hybridization

Fluorescent in situ hybridization experiments were performed using the RNAscope Multiplex Reagent Kit v2 assay (ACD Bio) according to the manufacturer’s instructions for fixed-frozen tissue. Probes used included mouse *Apoe* (ACD Bio #590131-C4; Lot #21270A, 23346D), *Abca1* (ACD Bio #522251; Lot #25072B), *Hexb* (ACD Bio #314231-C3; Lot #23346D, 22285B), and human *APOE* (ACD Bio #1044891-C1; Lot #21351B). For post-hybridization immunofluorescent labeling, glass-mounted sections were incubated with blocking and antibody solutions as described above. Sections were imaged on a Leica SP8 confocal microscope using a 63× objective (total z-depth, 12 µm; z-step size, 0.5 µm).

### RNAscope In situ Transcript Analysis

Fluorescent in situ hybridization images were analyzed using Imaris software (Oxford Instruments, v10.0.0). When Iba1 protein immunostaining was present, microglia were identified by constructing three-dimensional (3D) surfaces based on Iba1 signal. Three-dimensional surfaces corresponding to *Apoe* and *Abca1* transcripts were generated, and transcript volume overlapping with microglial surfaces was quantified and normalized to total microglial volume per image. In experiments in which *Hexb* transcript signal was used to identify microglia, 3D surfaces were generated based on *Hexb* expression. The volume of *Apoe* transcript signal overlapping with *Hexb* surfaces was quantified and expressed relative to total *Hexb* transcript volume per image.

### Real-Time Quantitative PCR

Liver tissue was perfused with sterile saline and rapidly frozen in liquid nitrogen and stored at −80C until RNA extraction. Total RNA was extracted using TRIzol, measured with NanoDrop (Thermo Scientific), and reverse-transcribed with *Power*SYBR Green PCR Master Mix (Applied Biosystems). Transcript levels were measured by semiquantitative PCR on ABI QuantStudio 5 Real-Time PCR system (Applied Biosystems). Relative mRNA abundance was normalized to 18S. Primers were designed using PrimerBank. See sequences below:

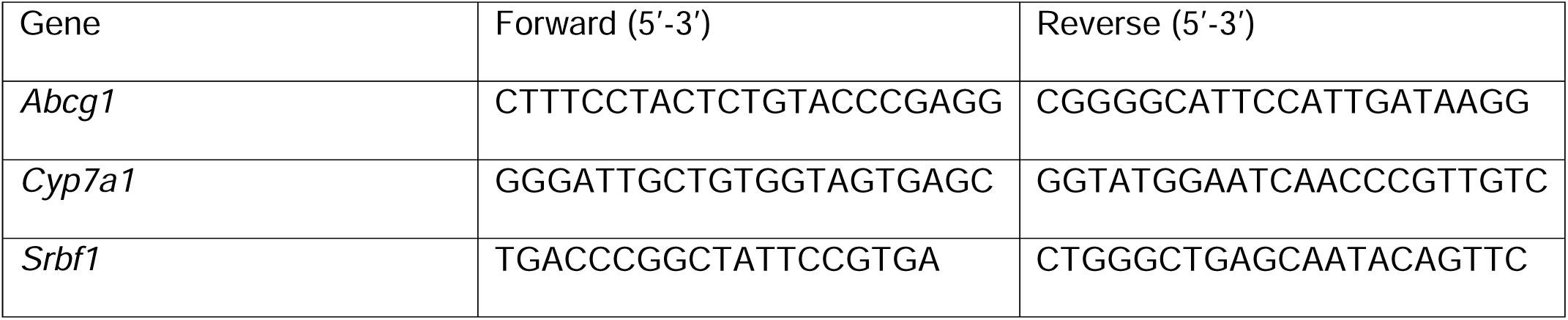

QUANTIFICATION AND STATISTICAL ANALYSIS

KEY RESOURCES TABLE

