## Supplemental Figure Legends for "ApoE-Dependent Lipid Handling by Median Eminence Microglia Preserves Myelin Integrity and Metabolic Function"

**Figure S1 (related to Figure 1). FACS gating strategy for microglial enrichment from the mediobasal hypothalamus.**

Representative flow cytometry plots show sequential gating to exclude debris and doublets, select viable cells ($DAPI^{-}$), and enrich for $CD11b^{+}$ myeloid cells from mediobasal hypothalamus (MBH) dissociates for downstream analyses.

**Figure S2 (related to Figure 1). Validation of MULTI-seq barcoding and sample representation across dietary conditions.**

(A) Barcoding efficiency and accuracy are shown by barcode validation plots, with the number of cells recovered for each experimental group indicated.

(B) UMAP projections display integrated single-cell transcriptomic profiles from mediobasal hypothalamic microglia of mice fed a control diet (CD) or western diet (WD).

(C) Quantification showing comparable representation of cell proportions across dietary conditions.

**Figure S3 (related to Figure 2). Transcriptional characterization of MG-Apoe subclusters.**

(A) Dot plot showing expression of selected marker genes across MG-Apoe subclusters (MG-$Apoe^{hi}$, MG-$Apoe^{int}$, and MG-$Apoe^{lo}$), with dot size indicating the fraction of cells expressing each gene and color intensity (blue to red) reflecting scaled log fold change.

(B) Mean disease-associated microglia (DAM) signature scores across MG-Apoe subclusters, cycling microglia, and homeostatic microglia.

(C) Mean interferon-responsive microglia (IRM) signature scores across the indicated microglial populations.

(D) Mean senescence-associated microglia (SenMayo) signature scores across the indicated microglial populations.

Data are presented as mean ± SEM.

**Figure S4 (related to Figure 3).** ***Itgax*** **(CD11c) mRNA is enriched in median eminence microglia and confined to myelinated regions.**

(A) Differential expression analysis showing enrichment of *Itgax* in MG-Apoe microglia compared with homeostatic microglia.

(B) Violin plots illustrating *Itgax* expression across MG-Apoe subclusters and homeostatic microglia.

(C) Representative 20× images demonstrating that CD11c expression is spatially restricted to the myelinated region of the median eminence (ME) and absent from adjacent hypothalamic nuclei, including the arcuate nucleus (ARC) and ventromedial hypothalamus (VMH). Scale bars, 50 μm.

(D) Quantification of $CD11c^{+}$ volume in the ME, ARC, and VMH.

(E) Quantification of myelin basic protein (MBP) signal showing increased myelin content in the ME relative to the ARC and VMH.

(F) Representative images showing increased CD68 and STAT1 immunoreactivity in the ME compared with adjacent hypothalamic regions. Scale bars, 50 µm.

(G) Quantification of CD68 signal intensity.

(H) Quantification of STAT1 signal intensity.

(I) Representative images demonstrating overlap between CD11c and Clec7a expression in ME microglia, indicating that both markers identify a shared activation program. Scale bars, 20 µm.

Data are presented as mean ± SEM. 3V, third ventricle.

**Figure S5 (related to Figure 3). Myelin production selectively supports Clec7a$^{+}$ microglial program but not interferon signaling in the median eminence.**

(A) Representative images showing myelin basic protein (MBP) immunoreactivity and the microglial marker Iba1 in the median eminence (ME) of adult male Myrf$^{fl/fl}$ control (OPC-Myrf $^{CTL}$) and Pdgfrα-CreER;Myrf conditional knockout (OPC-Myrf $^{KO}$) mice following tamoxifen-induced deletion and 4 weeks of western diet (WD) feeding.

(B) Quantification of total MBP volume in the ME.

(C) Quantification of MBP signal within Iba1$^{+}$ microglia, indicating reduced microglial myelin processing following Myrf deletion in oligodendrocyte progenitor cells.

(D) Representative images showing Clec7a expression in Iba1$^{+}$ microglia within the ME of OPC-Myrf $^{CTL}$ and OPC-Myrf $^{KO}$ mice during WD feeding.

(E) Quantification of Clec7a$^{+}$ microglia demonstrating attenuation of Clec7a induction following *Myrf* deletion.

(F) Quantification of STAT1$^{+}$ microglia demonstrating preserved interferon signaling despite impaired myelin production.

Data are presented as mean ± SEM. Statistical significance was assessed using unpaired two-tailed t-tests. Scale bars, 20 µm.

**Figure S6 (related to Figure S4). Western diet feeding induces Trem2 expression in median eminence microglia.**

(A) Representative images showing Trem2 immunoreactivity with the microglial marker Iba1 in the hypothalamic median eminence (ME) of adult male mice fed a control diet (CD) or western diet (WD) for 4 weeks.

(B) Quantification of Trem2$^{+}$ volume within Iba1$^{+}$ microglia in the ME, demonstrating increased Trem2 expression following WD feeding (n = 4–6 mice per group; *$p < 0.05$).

Data are presented as mean ± SEM. Statistical significance was assessed using unpaired two-tailed t-tests.

**Figure S7 (related to Figures S4 and S5). Body weight trajectories across genetic models during western diet feeding.**

(A) Body weight curves for *Apoe*$^{-}$/$^{-}$ mice and corresponding wild-type controls measured over 4 weeks of western diet (WD) feeding (n = 4–5 mice per group; **p < 0.01, ***p < 0.001).

(B) Body weight curves for *Trem2*$^{-}$/$^{-}$ mice and corresponding wild-type controls measured over 4 weeks of WD feeding (n = 10 mice per group; *p < 0.05).

(C) Body weight curves for humanized APOE3/E3 and APOE4/E4 mice measured over 4 weeks of WD feeding (n = 5 mice per group; *p < 0.05).

Data are presented as mean ± SEM. Statistical significance was assessed using unpaired two-tailed t-tests.

**Figure S8. Validation of sHDL–LXR nanoparticle uptake by median eminence microglia and assessment of peripheral lipogenic gene expression.**

(A) Representative images showing uptake of DiD-labeled sHDL nanoparticles in the hypothalamic median eminence (ME), with colocalization to Iba1$^{+}$ microglia and enrichment near the third ventricle (3V).

(B) Higher-magnification (60×) image confirming intracellular localization of DiD–sHDL nanoparticles within ME microglia.

(C) Quantitative PCR analysis of hepatic lipogenic and cholesterol-related genes (*Abcg1*, *Cyp7a1*, *Srebf1*) showing no significant changes following sHDL–LXRa treatment compared with sHDL-blank controls. Data are presented as mean ± SEM.

(D) Representative RNAscope images showing *Abca1* transcripts in Iba1$^{+}$ microglia within the ME of mice treated with sHDL-blank or sHDL–LXRa.

(E) Quantification of *Abca1* transcript signal within Iba1$^{+}$ microglia, demonstrating increased *Abca1* expression following sHDL–LXRa treatment.
