## Supplemental Figure 1 for "ApoE-Dependent Lipid Handling by Median Eminence Microglia Preserves Myelin Integrity and Metabolic Function"

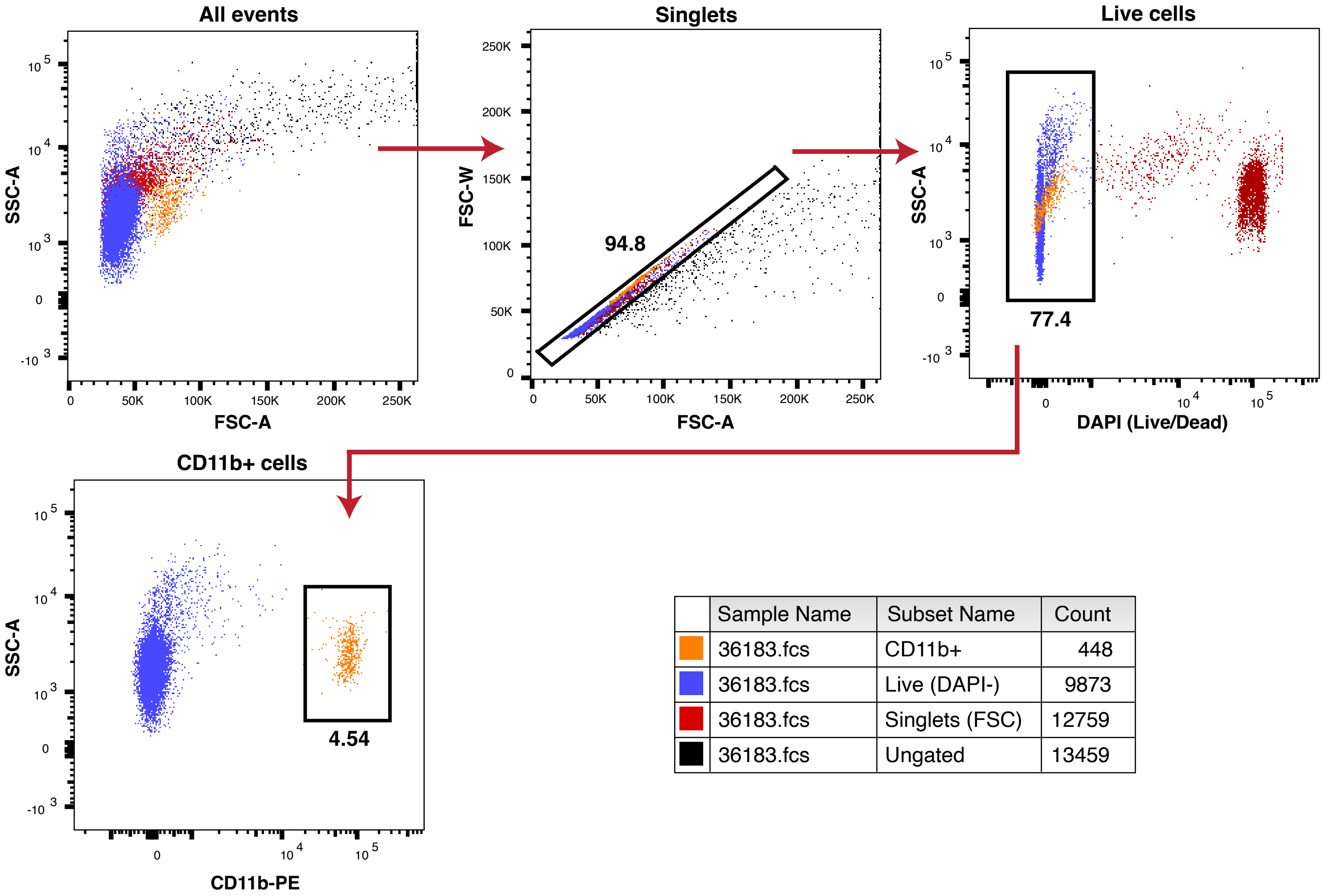


| | Sample Name | Subset Name | Count |
|---|---|---|---|
| (orange) | 36183.fcs | CD11b+ | 448 |
| (blue) | 36183.fcs | Live (DAPI-) | 9873 |
| (red) | 36183.fcs | Singlets (FSC) | 12759 |
| (black) | 36183.fcs | Ungated | 13459 |
