## Supplementary figures and images for "ApoE-Dependent Lipid Handling by Median Eminence Microglia Preserves Myelin Integrity and Metabolic Function"

### Supplemental Figure 2

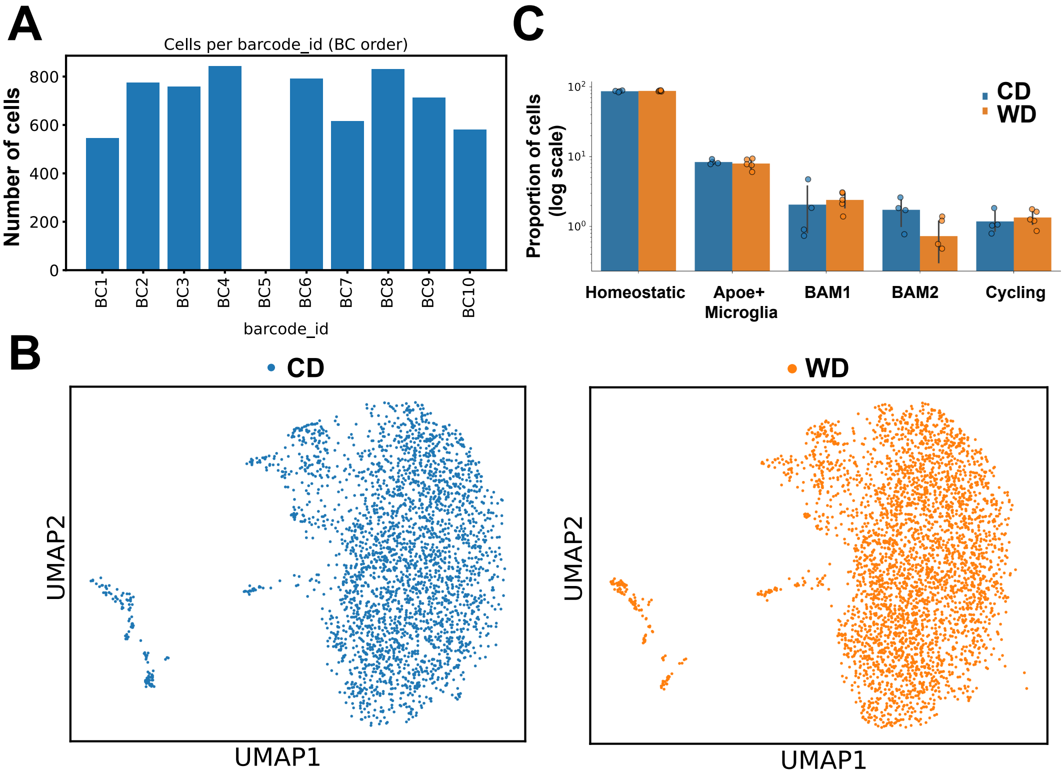

### Supplemental Figure 3

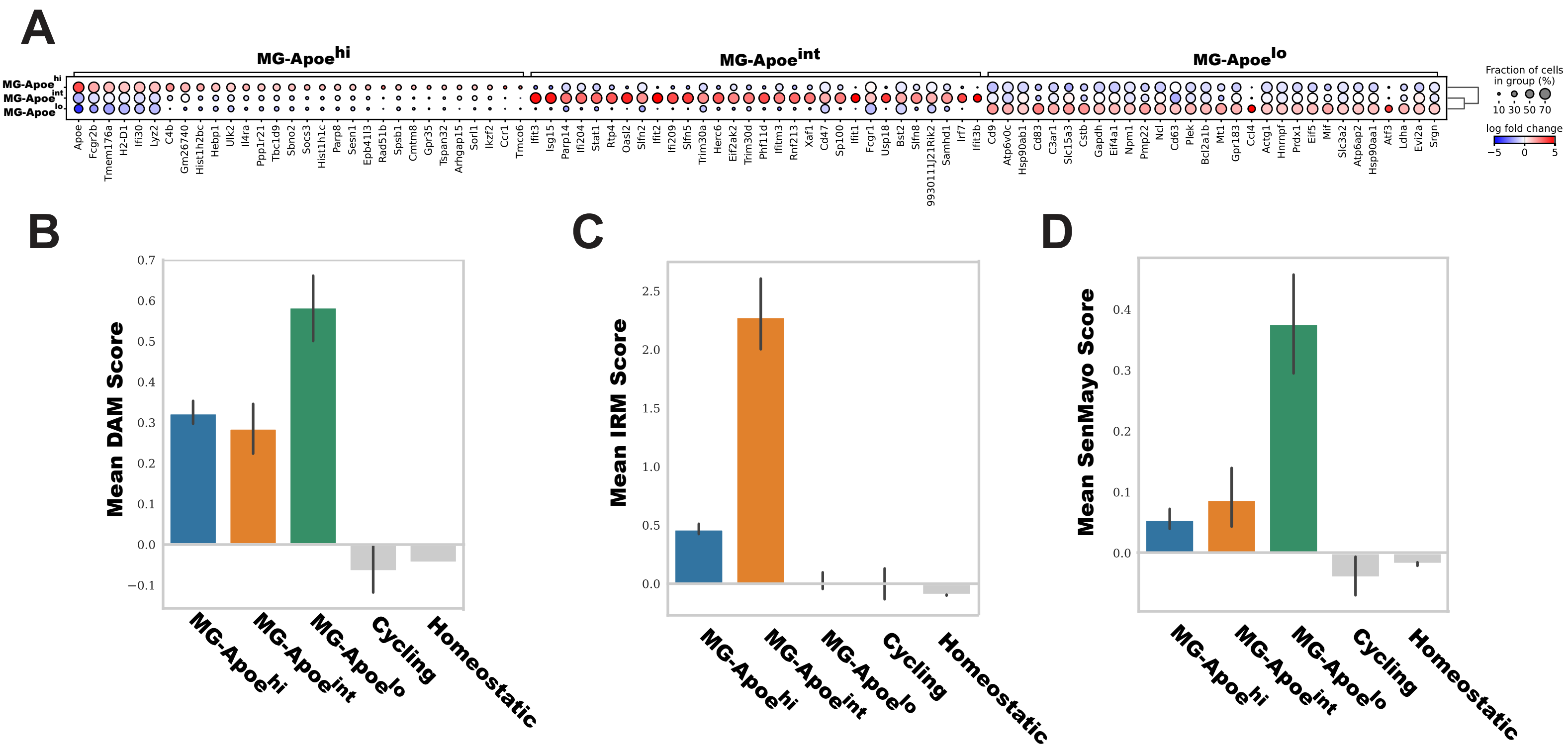

### Supplemental Figure 4

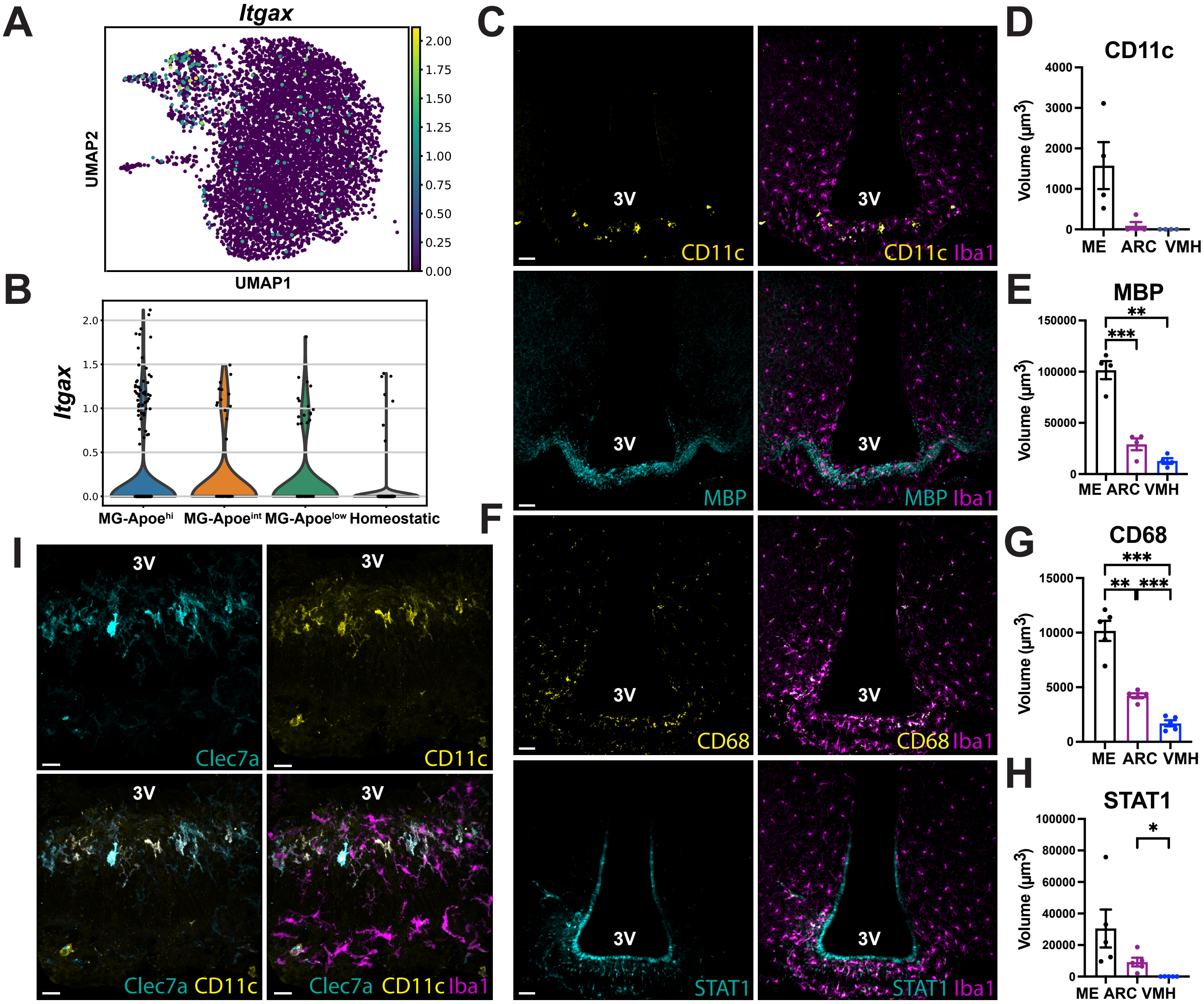

### Supplemental Figure 5

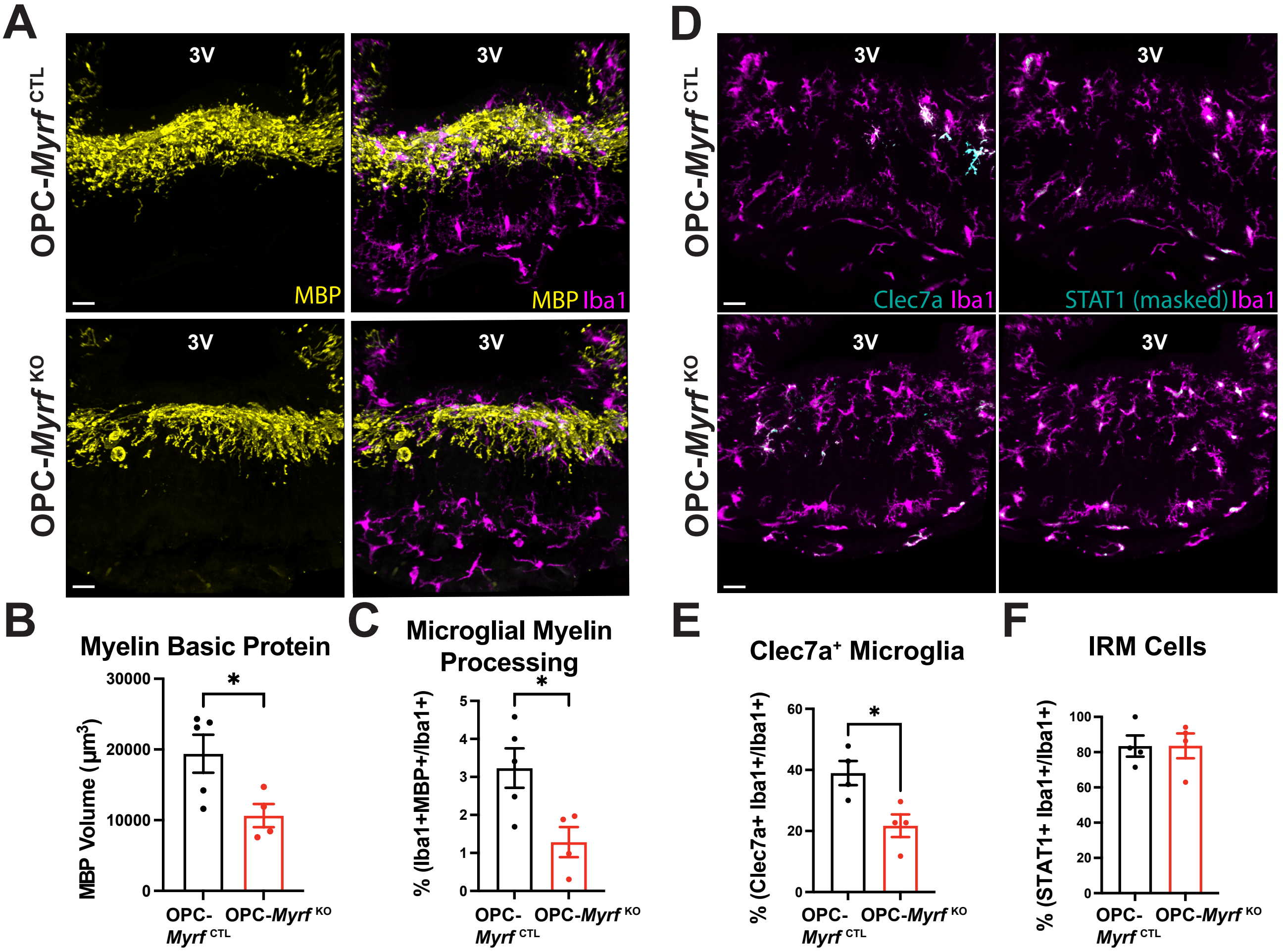

### Supplemental Figure 6

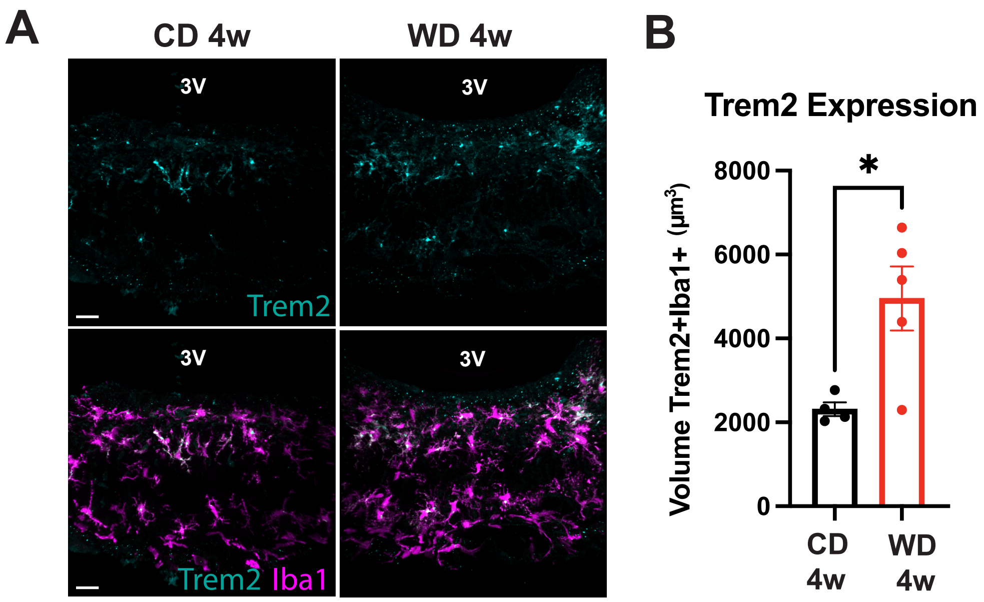

### Supplemental Figure 7

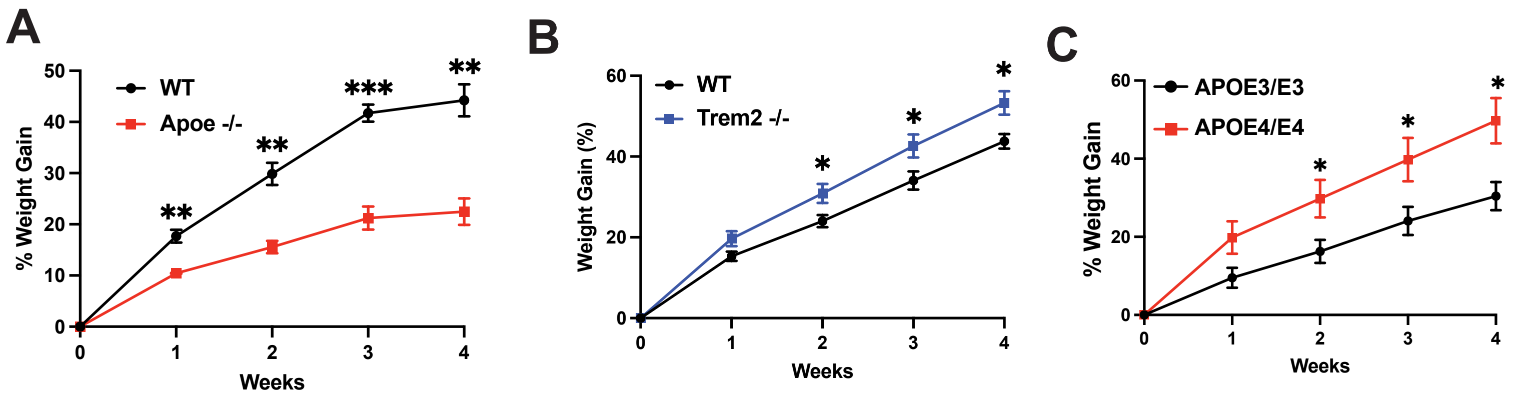

### Supplemental Figure 8

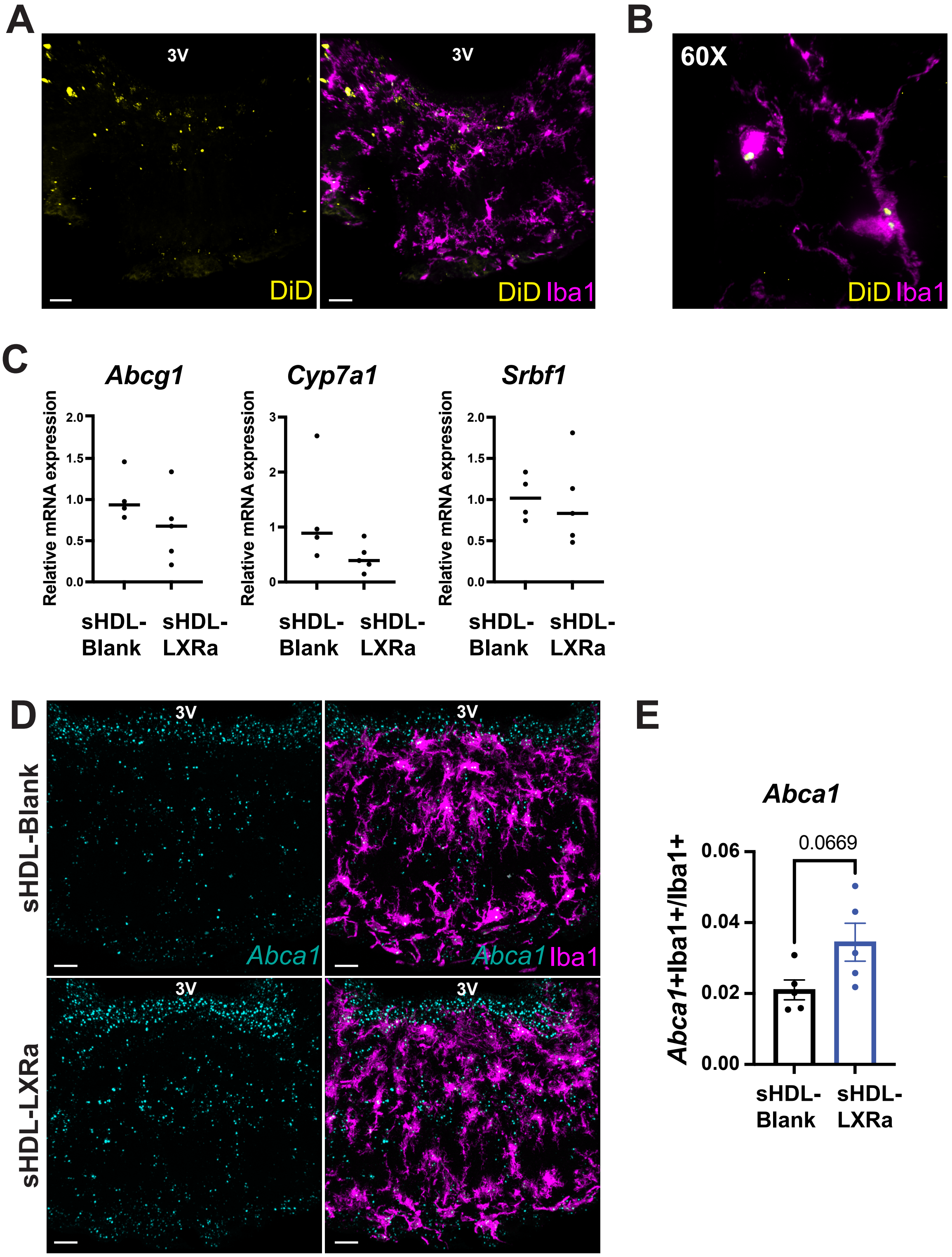
